# SARS-CoV-2 causes periodontal fibrosis by deregulating mitochondrial β-oxidation

**DOI:** 10.1101/2022.12.15.520561

**Authors:** Yan Gao, Wai Ling Kok, Vikram Sharma, Charlotte Sara Illsley, Sally Hanks, Christopher Tredwin, Bing Hu

**Affiliations:** Stem Cells & Regenerative Medicine Laboratory, Peninsula Dental School, Faculty of Health, University of Plymouth, 16 Research Way, Plymouth, PL6 8BU, UK; School of Biomedical Sciences, Faculty of Health, University of Plymouth, 16 Research Way, Plymouth, PL6 8BU, UK

**Keywords:** COVID-19, SARS-CoV-2, Tooth, Periodontal ligament, Fibrosis, Mitochondria

## Abstract

The global high prevalence of COVID-19 is a major challenge for health professionals and patients. SARS-CoV-2 virus mutate predominantly in the spike proteins, whilst the other key viral components remain stable. Previous studies have shown that the human oral cavity can potentially act as reservoir of the SARS-CoV-2 virus. COVID-19 can cause severe oral mucosa lesions and is likely to be connected with poor periodontal conditions. However, the consequence of SARS-CoV-2 viral infection on human oral health has not been systematically examined. In this research, we aimed to study the pathogenicity of SARS-CoV-2 viral components on human periodontal tissues and cells. We found that by exposing to SARS-CoV-2, especially to the viral envelope and membrane proteins, the human periodontal fibroblasts could develop fibrotic pathogenic phenotypes, including hyperproliferation that was concomitant induced together with increased apoptosis and senescence. The fibrotic degeneration was mediated by a down-regulation of mitochondrial β-oxidation in the fibroblasts. Fatty acid β-oxidation inhibitor, etomoxir treatment could mirror the same pathological consequence on the cells, similar to SARS-CoV-2 infection. Our results therefore provide novel mechanistic insights into how SARS-CoV-2 infection can affect human periodontal health at the cell and molecular level with potential new therapeutic targets for COVID-19 induced fibrosis.

## Introduction

Since its outbreak, the coronavirus disease 2019 (COVID-19) pandemic has infected over 63 million people globally (https://covid19.who.int) and has been a major challenge to human health. The severe acute respiratory syndrome coronavirus 2 (SARS-CoV-2 virus) has multiple transmission routes such as through respiratory fluids, therefore is highly infectious (Ferretti et al. 2020). It has been reported that there were ever increasing “long COVID” (Lopez-Leon et al. 2021) and repeated infection cases (Wang et al. 2021), with the UK alone already having more than 2 million long COVID cases (Wise 2022). Although COVID-19 has been initially connected with acute inflammatory disease, which causes progressive pulmonary fibrosis (Spagnolo et al. 2020), increasing evidence have suggested that COVID-19 does affect other organs such as heart, skin, kidneys and brain (Wang et al. 2020). The human oral cavity and saliva have also been demonstrated to be an important reservoir of the SARS-CoV-2 virus (Huang et al. 2021), and saliva have been used for effective diagnosis of COVID-19 (Baghizadeh Fini 2020). Periodontal tissues are highly vulnerable to different infectious diseases and COVID-19 has been reported to be potentially connected with poor periodontal health (Qi et al. 2022). Furthermore, severe oral mucosa lesions including detachment of oral epithelium with inflammation and fibrosis have been reported in the COVID-19 patients (Henin et al. 2022; Pandiar et al. 2022). However, the molecular mechanisms under the potential pathological consequence of SARS-CoV-2 infection on human oral health have not been systematically investigated.

The evolution of SARS-CoV-2 has generated different variants that are responsible for infection speeds and symptoms (https://www.who.int/activities/tracking-SARS-CoV-2-variants). SARS-CoV-2 has four different structural protein components: envelope, membrane, nucleocapsid and spike (Figure 1A) (Huang et al. 2020). The mutations of SARS-CoV-2 predominantly occur in the spike proteins, particularly the receptor-binding domain (RBD), whilst the other key viral components remain stable (Satarker and Nampoothiri 2020) (Harvey et al. 2021). A key step of SARS-CoV-2 infection is the binding of spike RBD to angiotensin-converting enzyme 2 (ACE2) receptor on the target cells (Ni et al. 2020) and co-receptor transmembrane serine protease 2 (TMPRSS2) to trigger viral internalization together with the molecular downstream cascades (Hoffmann et al. 2020). Currently, most of the COVID-19 related research and vaccine development have focused on the spike protein, leaving the other structural proteins’ pathological functions remain to be elucidated.

**Figure 1.**
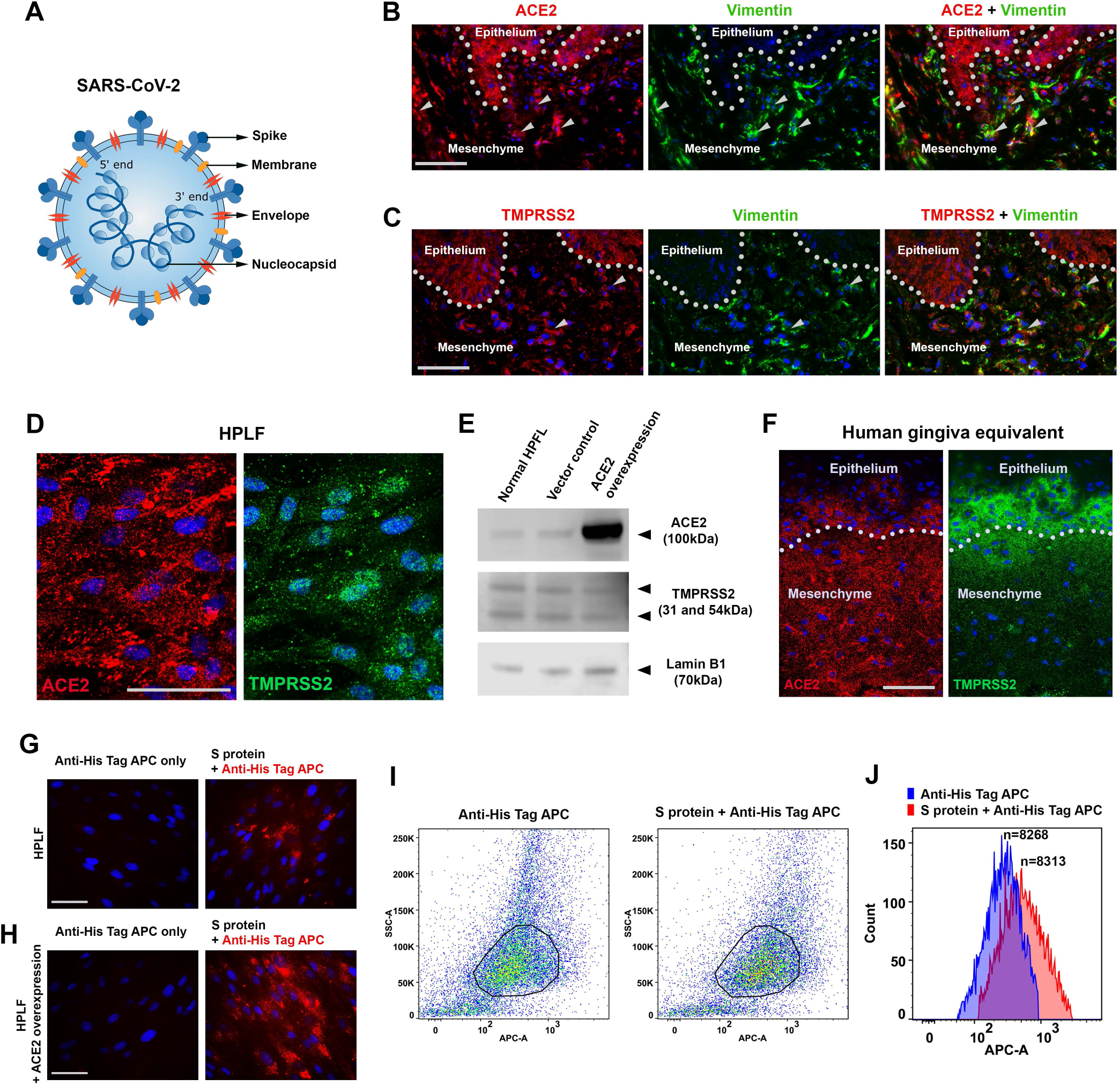
Human periodontal cells express ACE2 and TMPRSS2, and prone to SARS-CoV-2 infection. A. Illustration of SARS-CoV-2’s key structure proteins; B and C. Immunofluorescence analysis of ACE2 or TMPRSS2 co-expression with Vimentin in human periodontal tissues. using specific antibodies and Alexa 568 (red) or Alexa 488 (green) conjugated secondary antibodies. Dotted lines mark epithelia-mesenchymal junctions. Arrows indicate typical clusters of fibroblasts with co-expression of the indicated markers. D. Immunofluorescence analysis of ACE2 or TMPRSS2 in human periodontal ligament fibroblasts (HPLF). E. Western blotting analysis of ACE2 and TMPRSS2 expression in human periodontal ligament fibroblasts (HPLF) under normal growing condition, and lentiviral mediated ACE2 overexpression. Lamin B1 was used as loading control. All the blots were performed sequentially on the same membrane. F. Immunofluorescence analysis of ACE2 or TMPRSS2 in human gingiva equivalent (also see Appendix Figure 1A). G and H. Normal HPLF or the cells with ACE2 overexpression was treated with His tagged spike (S) protein first then traced using Anti-His Tag APC conjugated antibodies. For controls, the spike protein was omitted (also see Appendix Figure 1B). I and J: FACS analysis of the samples showed in E and F. Note the anti-His Tag APC antibody did show background but the shifting of the florescence peak could still be visualized (H). n: particle (cell) count. Bars: 100um

As part of a series of research on the biological connection of COVID-19 with oral health, this study intended to apply human periodontal tissue and cells as examples to dissect the direct pathological effects of SARS-CoV-2 viral structural components.

## Materials & methods

### Cell culture

Human periodontal ligament fibroblasts (HPLF) were cultured in DMEM/F12 ((Gibco, 31331-028) containing 20% Fetal bovine serum (FBS) (Sigma, F7524), 1% penicillin-streptomycin (Hyclone, SV30079.01). Human gingival epithelial cells (HGEPp) were cultured in CnT-57 (CELLnTEC, CnT-57).

### Viral infection

Plasmids for SARS-CoV-2 structural proteins were purchased from Addgene (Appendix Table 2). Lentiviral supernatant was collected according to the manual using 293FT cells. HPLFs were infected with lentiviruses carrying target sequences above with 10 μg/ml polybrene (Merck, TR-1003). After 2 h, dishes were topped up with fresh culture medium. Samples were collected at 6 h or 48 h. For overexpressing ACE2 (Addgene, Appendix Table 2) in HPLFs, the cells were infected with lentiviruses carrying target sequences above with 10 μg/ml polybrene (Merck). After 2 h, dishes were topped up with fresh normal culture medium. Samples were collected according to different time points. Infected cells were selected with 1 μg/ml puromycin (Thermo Scientific, 10781691) for 7-10 days.

### Recombinant SARS-CoV-2 spike protein treatment

Recombinant SARS-CoV-2 spike protein with His-tag (Biotechne, 10549-CV) was diluted with culture medium to 500 ng/ml or 5 μg/ml and added into the cell culture medium. Samples were collected at 6 h or 48 h.

### Mitochondria β-oxidation inhibition assay

Etomoxir (Sigma, E1905) was diluted with culture medium to aimed concentration then added into cell culture medium. Samples were collected at 6 h or 48 h.

### Human periodontal tissue 3D equivalents

5*10^4 of HPLFs were mixed with 150 μl gel mixture of rat tail collagen (Fisher, 11519816), DMEM (Fisher, 21969-035) and FBS (Ratio of volume 9:1:1) homogenously on ice. 2.3 μl 1M Sodium hydroxide solution (NaOH, Sigma 71687) was added into the mixture for neutralization, 150 μl gel was pipetted into a 0.4 μm culture insert (Greiner, 662641) incubated at 37°C for 1 h then fresh culture medium was added into the insert. Culture medium was replaced by CnT-57 before seeding 5*10^5 of HGEPp on top of the gel. HGEPp was cultured for 72 h before airlifting. Culture medium inside the insert was removed every day and the medium outside was changed every 2 days. Samples were frozen directly in OCT (Agar Scientific, AGR1180) at day 14 and sectioned at 20 μm for further analysis.

### Hydrogel based 3D matrix production assay

1*10^6 of HPLFs were mixed with 100 μl bioink (CELLINK, CELLINK SKIN+) slowly and gently. Gels were pipetted into 6 well plate then 1.5 ml crosslinking agent (CELLINK) was added to cover the gel at RT for 5 min. Crosslinking agent was removed and 1.5 ml fresh culture media was added which was replaced every 2 days. Samples were frozen directly in OCT at day 7 and sectioned at 30 μm for further analysis.

### Immunohistochemistry

For the details of immunostaining, BrdU incorporation assay, Terminal deoxynucleotidyl transferase dUTP nick end labelling (TUNEL) assay and Senescence assay details please see **Appendix materials & methods.**

### Flow cytometry analysis

HPLFs and ACE2 overexpressed HPLFs were harvested and fixed in 2% PFA solution in 10 mM PBS for 10 min then washed with FACS buffer (1% BSA in PBS). 1*10^6 cells were resuspended in 500 μl flow cytometry permeabilization buffer (0.1% Tween-20 in PBS) for 15 min then washed with FACS buffer again. Cells were resuspended in 100 μl FACS buffer and APC His-Tag conjugated antibody was added, incubated for 2 h at room temperature then kept in dark at 4 °C degree overnight. Samples were analyzed using the BD FACSAria™ II (BD Biosciences). Data was acquired using red laser (633-640nm) for APC signal. Results were analyzed using the FlowJo software (Tree Star Inc., Version 10.8.1). Gates and regions were placed around populations of cells with common characteristics based on SSC and APC.

### Western blotting

A NuPage® Electrophoresis System (Thermo Fisher Scientific), 4-12% Bis-Tris gradient gel (Thermo Fisher Scientific, NP0335BOX), and MOPS buffer (Thermo Fisher Scientific, NP0001), 25-40 µg protein were used for protein separation. Transfer of protein samples onto a 0.45 µm PVDF membrane (Thermo Fisher Scientific, LC2005) was carried out using a NuPage® XCell II Blot Module, and transfer buffer (Thermo Fisher Scientific, NP0006) with 10% methanol (Sigma, 322415). The iBind™ Western System (Thermo Fisher Scientific) was used for blocking, primary and secondary antibody incubations which details could be found above. A C-Digit scanner (LI-COR) was used for band detection with Image studio software (LI-COR, Version 3.1).

### Proteomic analysis

Sample preparation, in-gel digestion, sample cleanup and mass spectrometric analysis was carried out as described previously (Dunn, J et al, 2018). For details please see **Appendix material & methods**.

### Real-time PCR and data analysis

Real-time RT-PCR analysis was performed on a LightCycler 480 Real-Time system (Roche) for 45 cycles, using a SYBR Green I MasterMix (Roche, 04887352001) and primers. 36b4 gene was used as housekeeping gene. Analyses were performed using three technical replicates using the 2-ΔΔCt method.

### Seahorse Mito Stress Test

3*10^3 of HPLFs per well were seeded into Seahorse XFe96 well plate (Agilent Technologies, 200941) for overnight. After 6 h and 48 h of viral infection, the medium was removed but a nominal 20 μl per well was left. Each well was washed twice with pre-warmed assay medium (Agilent Technologies, 103680). 80μl assay medium were added into each well, and cells were then incubated for 1 h at 37°C. Meanwhile, effector working solutions were prepared and loaded into ports of the XFe96 cartridge which was already rehydrated in XF buffer at 37°C overnight. The cartridge and the utility plate were inserted into XFe96 instrument to calibrate probes. Once the calibration was finished, the utility plate was replaced by cell plate and continued with the assay as indicated. When the progress was completed, cell plate was removed from the instrument. Medium was aspirated slowly from wells and all wells were gently washed with warmed assay buffer, Plates were stored at -20°C or continued with DNA assay for further normalization. Data were analyzed by Seahorse Analytics website and Wave software (Agilent Technologies, Version 2.6.3).

### Statistics

Statistical analyses were performed using Prism software (GraphPad software, Version 9.4.1). Unpaired t-test was applied to all measurements. Data from Seahorse mito stress assay was analyzed by Seahorse Analytics website and Wave software (Agilent Technologies, Version 2.6.3). All quantification and real time RT-PCR results were presented using style of Mean and Standard Deviation (error bars). Observed differences were calculated for p-values: * p<0.05; ** p<0.01; ***: p<0.001; ****: p<0.0001.

## Results

### Human periodontal tissues and fibroblasts express SARS-CoV-2 receptors

To achieve robust evidence of the presence of SARS-CoV-2 receptors: ACE2 and TMPRSS2 in human periodontal tissue and cells, using immunofluorescence analysis, we confirmed that both ACE2 and TMPRSS2 were notably expressed in the gingiva epithelium, as well as in the periodontal ligament (PDL) cells (Figure 1B and C). By analyzing the cultured HPLF, we could also confirm the presence of the two receptors by immunofluorescence (Figure 1D) and western blotting analysis in those cells (Figure 1E). With an established collagen gel based 3D human gingiva tissue equivalent using gingival epithelial cells and HPLF (Appendix Figure 1A), we also observed consistent clear expression of ACE2 and TMPRSS2 both in the epithelial cells and PDL fibroblasts (Figure 1F).

### SARS-CoV-2 has high affinity to PDL fibroblasts

We next explored the potentiality of SARS-CoV-2 in infecting PDL fibroblasts. By treating the cells with His Tag conjugated spike protein, followed by immunofluorescent analysis of the spike protein location using an anti-His Tag APC conjugated antibody (Appendix Figure 1B), we could observe clear association of the spike protein with the cells either under natural growing condition (Figure 1G), or in lentiviral mediated ACE2 overexpression condition (Figure 1H). The affinity of spike protein to the cells could be further validated by fluorescence activated cell sorting (FACS) analysis (Figure 1I and J). Therefore, we confirmed that SARS-CoV-2 indeed could infect PDL fibroblasts directly.

### SARS-CoV-2 envelope and membrane proteins induce fibrotic pathogenic phenotypes in PDL fibroblasts

A key pathological hallmark of COVID-19 infection is fibrosis in the lung and potentially also in the other tissues (Spagnolo et al. 2020). To evaluate the consequence of COVID-19 infection on the PDL, we either adopted lentiviral mediated SARS-CoV-2 infection for envelope, membrane or nucleocapsid proteins (Gordon et al. 2020), or applied recombinant spike proteins at 500ng/ml or 5ug/ml, to cultured human PDL fibroblasts. We simulated acute and long infection by infecting or treating PDLs for 6 hours or 48 hours. Cell proliferation was evaluated using Bromodeoxyuridine (BrdU) incorporation followed by immunostaining for anti-BrdU antibodies. The results indicated that under our tested conditions, only membrane protein significantly increased cell proliferation at 6 hours, while at 48 hours, the envelope and membrane proteins could both elevate the BrdU incorporation index (Figure 2A and B; Appendix Figure 2A). Spike protein, on the contrary, could not modulate cell proliferation instead (Figure 2C and D; Appendix Figure 2B).

**Figure 2.**
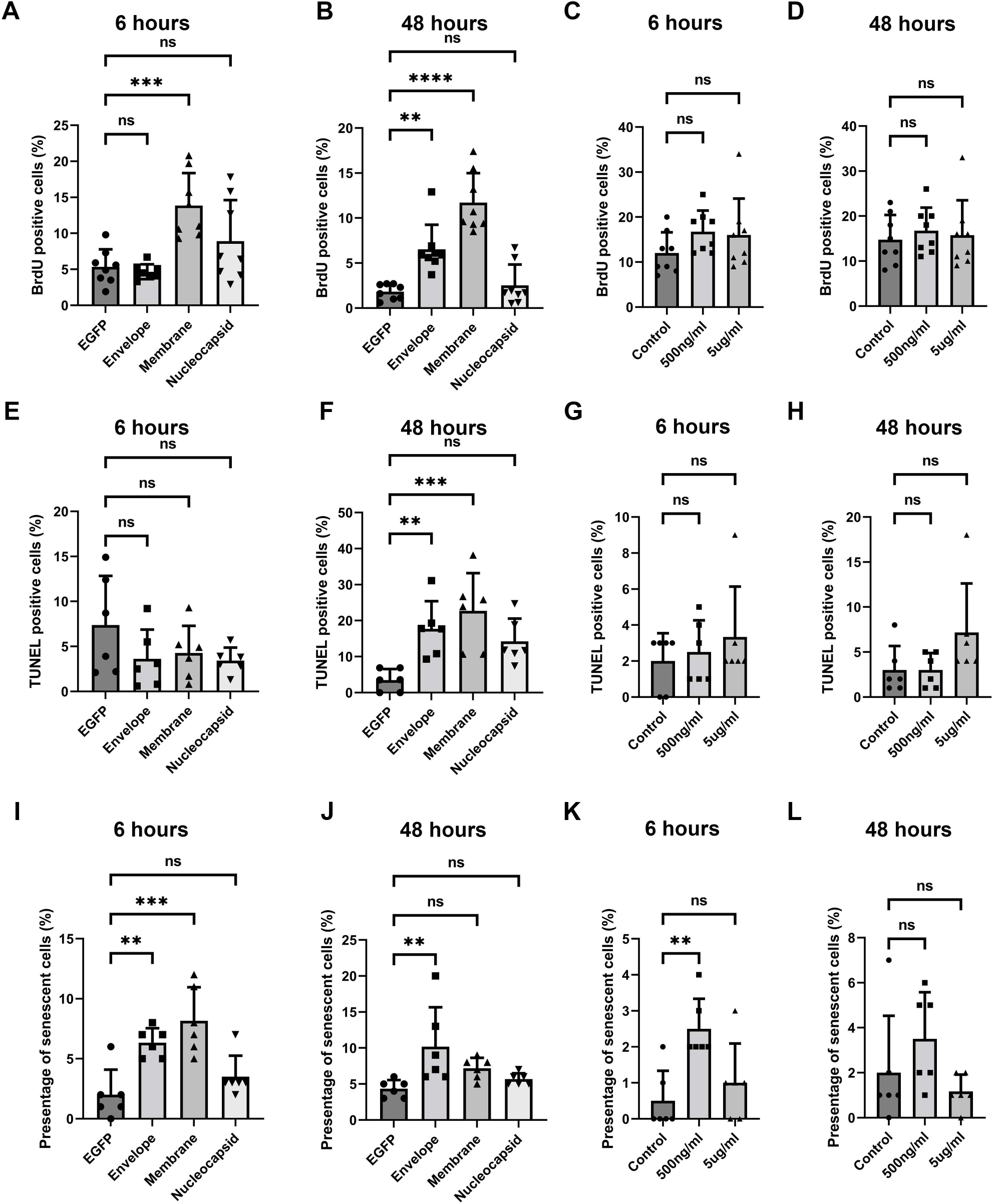
SARS-CoV-2 envelope and membrane protein could induce PDL fibroblast proliferation, apoptosis and senescence. A-D. BrdU incorporation analysis on the PDL fibroblasts treated with indicated conditions and time. Each data dot represent one random field in triplicated samples. Dunnett’s test was used for statistical analysis. Representative images can be found from Appendix Figure 2. E-H. TUNEL analysis on the PDL fibroblasts treated with indicated conditions and time. Each data dot represent one random field in triplicated samples. Dunnett’s correction was used for statistical analysis. Representative images can be found from Appendix Figure 3. I-L. Quantification of senescence-associated β-galactosidase positive cells (for identifying senescence) images on the indicated conditions. Each data dot represent one random field in triplicated samples. Representative images can be found from Appendix Figure 4. ns: no significance; ** p<0.01; ***: p<0.001.

With Terminal deoxynucleotidyl transferase dUTP nick end labelling (TUNEL), we then evaluated apoptosis status in the cells. The results indicated that at 48 hours, again only the envelope and membrane proteins, but not the nucleocapsid nor spike proteins could increase apoptosis (Figure 2E-H; Appendix 3A and B). In the meanwhile, for cellular senescence status in the tested conditions, the results showed that at 6 and 48 hours, only the envelope and membrane protein groups resulted in a significant increase in senescent cells (Figure 2I-L; Appendix 4A and B).

We then investigated extracellular matrix production by focusing on Collagen I and MMP1, the two key components and enzymes responsible for PDL tissue integrity. Interestingly, western blotting analysis revealed that all the SARS-CoV-2 structure proteins could induce Collagen I production (Figure 3A-F), and reduce MMP1 production, spike protein elevated MMP1 expression under one of the two tested doses (Figure 3A-F). Real time RT-PCR analysis suggested the Collagen I expression induction and MMP1 downregulation were modulated at transcription levels (Figure 3G and H). By applying a hydrogel based 3D culture system, we evaluated Collagen I and MMP1 production and deposition at three dimensions. The results showed that Collagen I deposition were again highly elevated, especially in the envelope and membrane protein groups (Figure 3I and J). And membrane and nucleocapsid proteins could bring down the MMP1 expression (Figure 3K and L).

**Figure 3.**
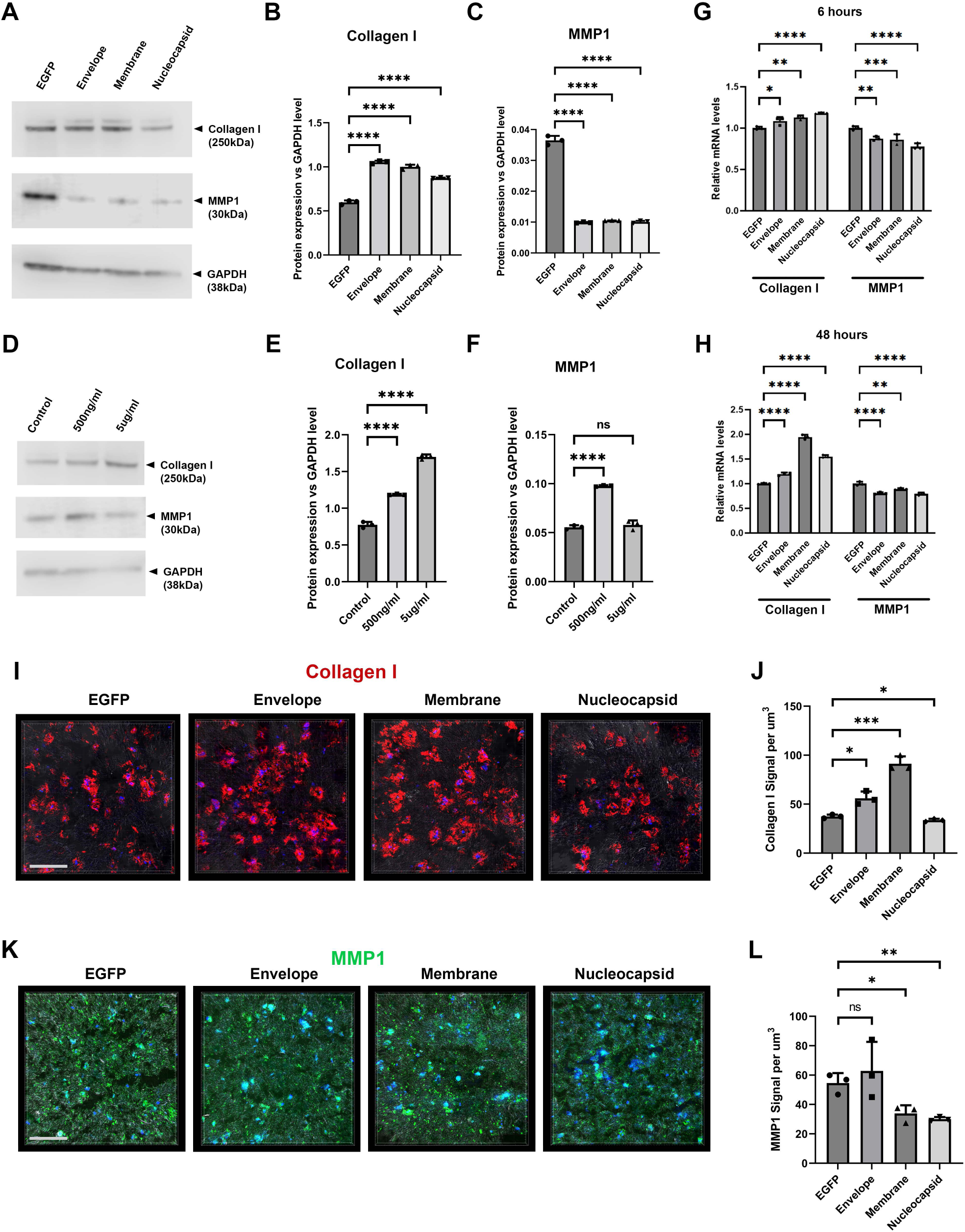
SARS-CoV-2 infection could induce collagen matrix deposition and MMP1 reduction. A-F. Western blotting analysis of Collagen I and MMP1 production in the cells under indicated conditions. A shows blotting results. B and C represents statistical analysis (each dot represents a single individual measurement). Dunnett’s test was used for statistical analysis. ns: no significance; G and H. Real time RT-PCR analysis of Collagen 1 and MMP1 transcription in the indicated conditions. I-L. Representative three dimensional analysis of Collagen I and MMP1 expression in the indicated conditions at 7 days after seeding the cells and infection in the hydrogel. The gel were stained using anti-Collagen I and MMP1 specific antibodies and developed with Alexa 568 (red) or 488 (Green) conjugated secondary antibodies. Each data dot represent one random field in triplicated samples. ns: no significance; * p<0.05; ** p<0.01; ***: p<0.001; ****: p<0.0001. Bars: 100um

Together, with the evidence of elevated fibroblast proliferation, apoptosis and senescence concomitantly, alongside increased matrix deposition and reduced metalloprotease production, our results pointed out that the SARS-CoV-2’s envelope and membrane were the most potent components to induce a fibrotic degeneration phenotype in PDL cells.

### SARS-CoV-2 envelope and membrane proteins down-regulate mitochondrial β- oxidation

To understand the molecular regulation mechanisms behind the observed pathological consequence of SARS-CoV-2 infection, we performed proteomic analysis on the treated PDL fibroblasts, by focusing on the effects of the envelope, membrane and nucleocapsid proteins. We then identified a group of proteins that were both suppressed at 6 and 48 hours post infection (Figure 4A; Appendix Table 1). Among the significantly down-regulated proteins, the trifunctional enzyme subunit alpha, isoform 2 of very long-chain specific acyl-CoA dehydrogenase, cytochrome c oxidase subunit 2 and isoform cytoplasmic of fumarate hydratase are all essential enzymes in mitochondria functions and mitochondrial β-oxidation (Figure 4A). Particularly the trifunctional enzyme subunit alpha (HADHA), isoform 2 of very long-chain specific acyl-CoA dehydrogenase were connected with mitochondrial fatty acid β-oxidation. In addition, western blotting analysis could further confirm that HADHA was indeed down-regulated by SARS-CoV-2 structural protein infection (Figure 4B-E).

**Figure 4.**
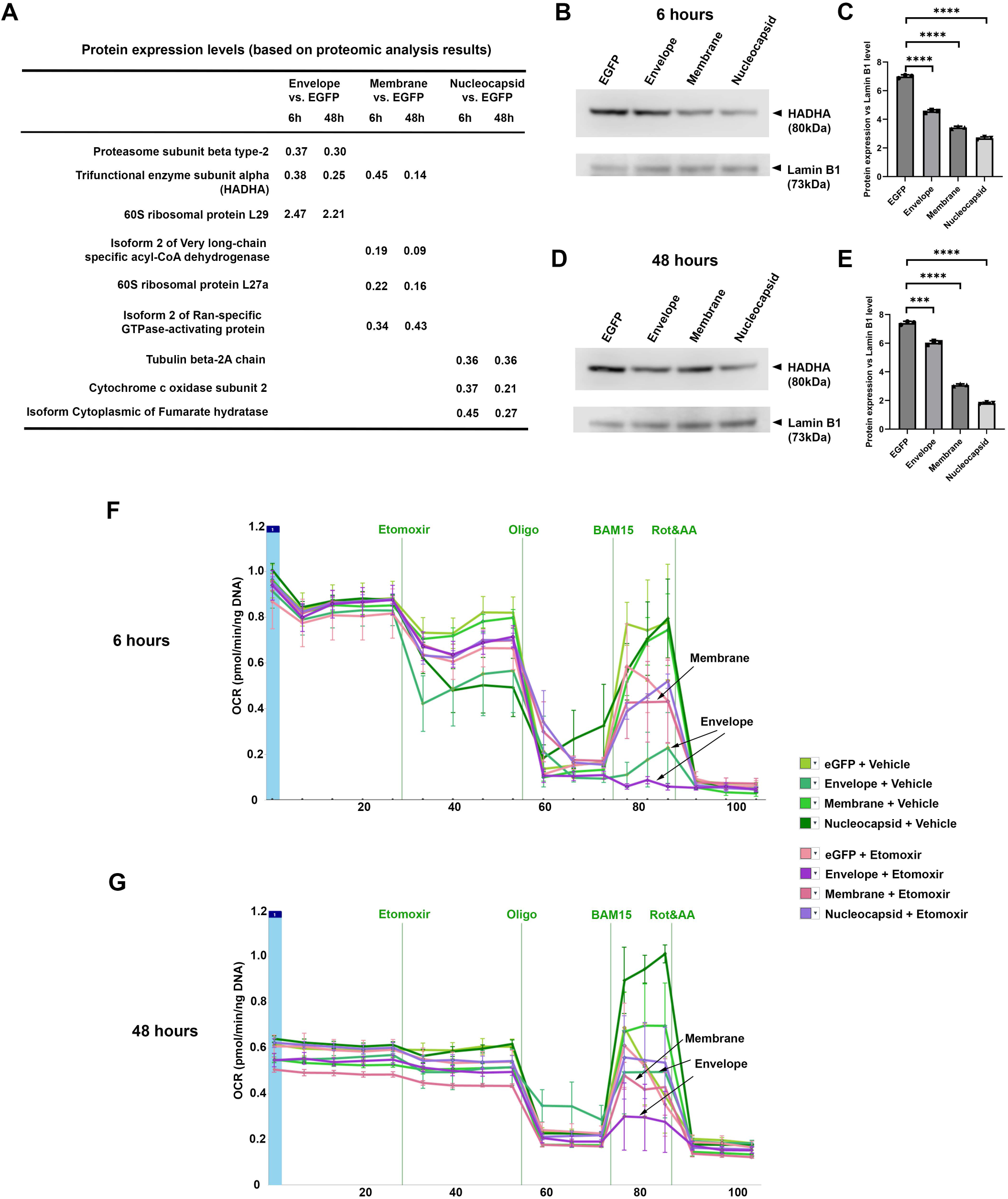
Envelope and membrane proteins are responsible to SARS-CoV-2 infection to PDL fibroblasts caused deregulate of mitochondria fatty acid β-oxidation pathway. A. Summary of proteomic analysis results of PDL fibroblasts treated with indicated conditions and time points. Only proteins showed <0.50 or >1.5 fold changes were included. For full data analysis please see Appendix Table 1. B-E. Western blotting analysis of HADHA expression in HPLF treated under indicated viral infection conditions either for 6 hours (B and C), or 48 hours (D and E). C and E represent statistical analysis (each dot represents a single individual measurement). ***: p<0.001; ****: p<0.0001. Lamin B1 was used as loading control. All the blots were performed sequentially on the same membrane. F and G. 6 and 48 hours Seahorse palmitate oxidation stress tests under indicated conditions either with vehicle alone or Etomoxir. Note for both time points, envelope group were down- regulated with/without etomoxir treatments, while membrane group also showed down regulation but only in the etomoxir treated samples.

We therefore conducted Seahorse Mito stress test (Appendix Figure 5) for evaluating if and how the mitochondria fatty acid pathway’s function could be affected by SARS-CoV-2 infection. The results showed that indeed, both SARS-CoV-2 envelope and membrane protein, could inhibit fatty acid β-oxidation at both 6 hours and 48 hours’ time points (Figure 4F and G).

**Figure 5.**
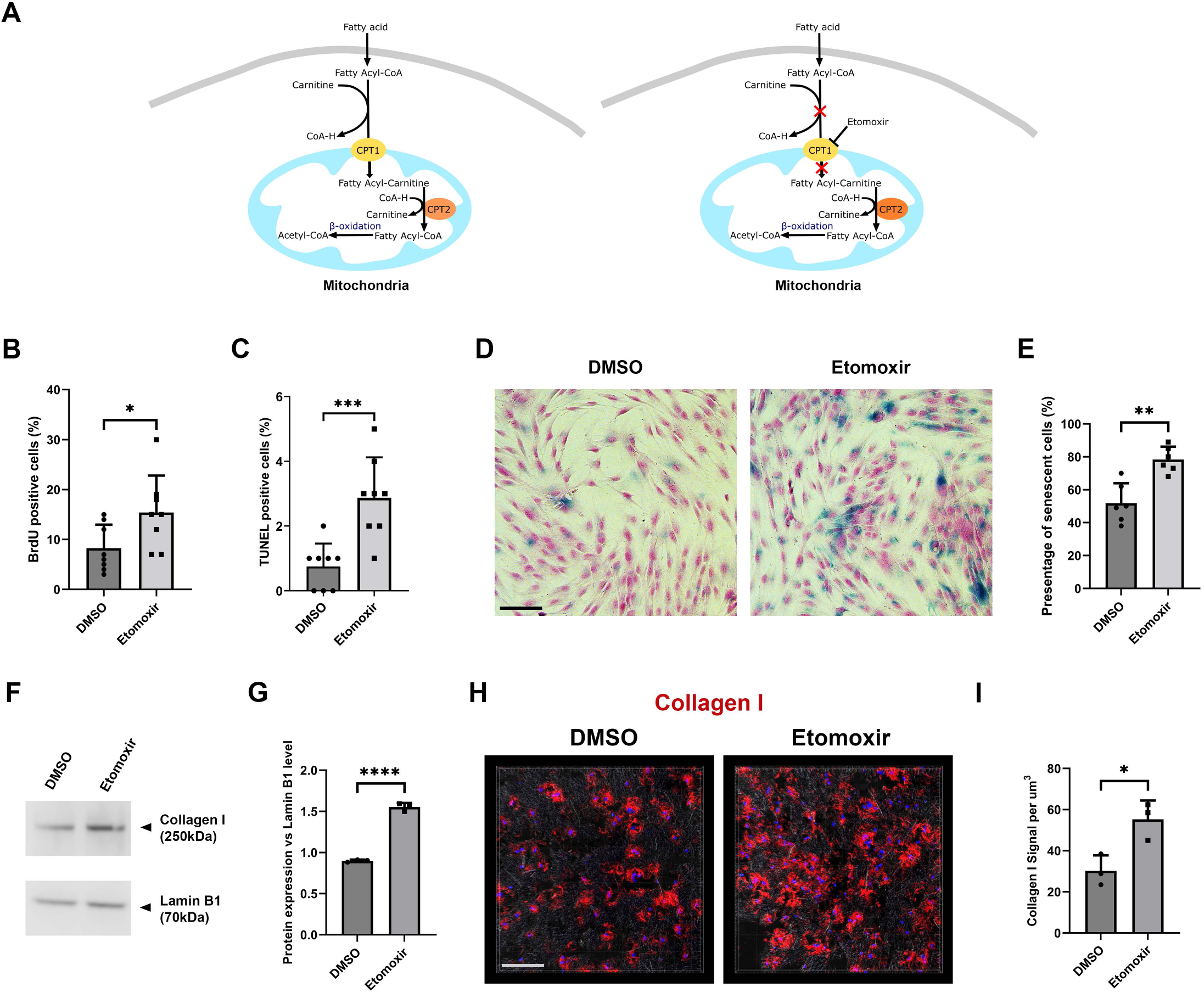
Mitochondrial fatty acid β-oxidation inhibition mirrored fibrotic degeneration phenotypes in PDL fibroblasts. A. Schematic drawing of fatty acid β-oxidation pathway (left) and the Etomoxir functioning mechanism (right). B and C. BrdU and TUNEL analysis of etomoxir treated cells. For representative images please see Appendix Figure 6. Each data dot represent one random field in triplicated samples. D and E. Representative senescence-associated β-galactosidase assay on indicated conditions. Note increased senescence in the etomoxir treated samples. Each data dot represent one random field in triplicated samples. F and G. Western blotting analysis of Collagen I expression in HPLF under etomoxir treatment or vehicle (DMSO) alone. Lamin B1 was used as loading control. All the blots were performed sequentially on the same membrane. G represents statistical analysis (each dot represents a single individual measurement). H and I. Representative 3D analysis of Collagen I expression in the indicated conditions at 48 hours after seeding the cells and etomoxir treatment in the hydrogel. The gel was stained using anti-Collagen I specific antibodies and developed with Alexa 568 (red) conjugated secondary antibodies. Each data dot represent one random field in triplicated samples. * p<0.05; ** p<0.01; ***: p<0.001; ****: p<0.0001. Bars: 100um

### Chemical inhibition of mitochondrial β-oxidation can mirror the fibrotic pathological consequence of SARS-CoV-2 infection

To validate the effect of fatty acid β-oxidation pathway inhibition, we next treated the cells with etomoxir, a specific inhibitor of the pathway through inhibiting carnitine palmitoyltransferase I (CPT1), a key regulatory enzyme for fatty acid to be imported into mitochondria (Figure 5A). The results indicated that etomoxir treatment could indeed mirror the same pathological consequence on the fibroblasts, similar to SARS-CoV-2 infection. We again observed significantly increased cell proliferation (Figure 5B; Appendix figure 6), apoptosis (Figure 5C; Appendix figure 6), and senescence (Figure 5D and E), together with elevated Collagen I production (Figure 5F and G). Similarly, in the hydrogel based three dimension PDL fibroblasts culture, Collagen I deposition was highly increased in the etomoxir treated samples (Figure 5H and I).

## Discussion

COVID-19 infection can cause a series of symptoms. Among them, fibrosis particularly in lung tissues has evoked significant attention due to the severe consequence on patient life and health quality. Although the current dominant SARS-CoV-2 variants (such as the Omicron) induce milder symptoms in human bodies, the exact pathological consequence of COVID-19 to different human tissues and organs, particularly for oral cavity tissues are still missing, possibly due to the difficulty of distinguishing conventional periodontal diseases from COVID-19 connections. Current concepts of COVID-19 etiology suggest lung fibrosis caused by SARS-CoV-2 infection can be mainly due to damages on lung epithelial cells that trigger acute inflammation followed by fibroblast hyperproliferation (Merad and Martin 2020). However, as SARS-CoV-2 infection is rapid and it is difficult to distinguish if the deeper cells (such as fibroblasts) can also be infected directly, such as in oral cavity the epithelial barrier could be rapidly disrupted upon infection (Henin et al. 2022) . We therefore cannot neglect the potential direct infection consequence of SARS-CoV-2 on fibroblasts. In particular, PDL is one of the most vulnerable human tissues that extrinsic virus and bacteria can often enter PDL directly in pathological conditions (Kononen et al. 2019). Consistent with this concept, previous reports have shown that ACE2 expression were elevated in the periodontal fibroblasts under gingivitis and periodontitis conditions(Santos et al. 2015), and could be induced by lipopolysaccharide, inflammatory cytokines etc. (Sena et al. 2021). As such, it is reasonable to postulate that SARS-CoV-2 can also directly infect PDL fibroblasts in the already damaged PDL, and can induce further pathological changes or enhance existing periodontal lesions. Attentions therefore should be made by dental clinicians to the patients who got COVID-19 and appeared to be diagnosed with periodontal disease at the same time, particularly for those long COVID and repeated infected cases.

Our results also suggested SARS-CoV-2 infection indeed can directly induce fibrotic disease phenotypes in fibroblasts through distinct pathways. Mitochondrial fatty acid β- oxidation is the major pathway responsible for fatty acids degradation, hence is essential for human body energy homeostasis. Impeding the pathway can cause different disorders (Merritt et al. 2018). Very recent studies have observed mitochondrial dysregulation in COVID-19 patient blood cells (Ajaz et al. 2021; Guntur et al. 2022). Interestingly, the dysfunction of the fatty acid oxidation has been previously connected with fibrosis particularly in the lung and kidney (Geng et al. 2021; Jang et al. 2020){Merritt, 2018 #12}. For the first time, our findings further confirmed that in the fibroblasts, SARS-CoV-2 could induce fibrotic degeneration directly, through down-regulating fatty acid β-oxidation, particularly by the virus’ envelope and membrane proteins.

Among the SARS-CoV-2’s structural proteins: envelope, membrane, nucleocapsid and spike, the first three proteins are stable structure proteins for all the reported variants, while so far all of the identified mutations happen inside the spike proteins (Harvey et al. 2021). Although spike protein is still the target especially for COVID-19 vaccine development, increasing evidence suggest the other SARS-CoV-2 structural proteins might have unexpected important roles in inducing COVID-19 symptoms. Previous structural analysis of the envelope protein suggested it might be important for virus pathogenicity (Mandala et al. 2020). The envelope protein can also physically increase intra-Golgi pH and forms cation channel(Cabrera-Garcia et al. 2021), and biochemically modulate spike protein in the meantime (Boson et al. 2021). The most abundant protein: the membrane protein in the SARS-CoV-2 virus, is rationally important for virus assembly (Zhang et al. 2022). Our results further confirmed that the envelope and membrane proteins are actually responsible for the fibrosis phenotypes in the cells by down regulating the key fatty acid β-oxidation regulator such as the trifunctional enzyme subunit alpha. The molecular mechanism behind this regulation axis would require further biochemical analysis.

In this study, our results provide novel mechanistic insights into how SARS-CoV-2 infection can affect human health, particularly for inducing fibrosis, at the cell and molecular level. The findings could be possibly extended to the other body systems to explain and explore the fibrosis pathology and treatment.

## Acknowledgement

We would like to thank the helps provided by Dr Jane Carré for assisting Seahorse analysis, Dr Paul Waines for FACS operation, and Prof. Simon Whawell for critical reading. This study was supported by a research grant from the European Orthodontic Society to Bing Hu. Yan Gao received a fellowship from the Peninsula Dental Social Enterprise.

## Author contributions

Yan Gao: Contributed to conception and design, acquisition, analysis, and interpretation, critically revised the manuscript.

Wai Ling Kok: Contributed to acquisition and analysis and critically revised the manuscript.

Vikram Sharma: Contributed to acquisition and analysis and critically revised the manuscript.

Charlotte Sara Illsley: Contributed to conception and design, and critically revised the manuscript.

Sally Hanks: Contributed to conception and design, and critically revised the manuscript.

Christopher Tredwin: Contributed to conception and design, and critically revised the manuscript.

Bing Hu: Contributed to conception and design, interpretation, and drafted and critically revised the manuscript.

All authors gave their final approval and agree to be accountable for all aspects of the work.

The authors declare that there is no conflict of interest regarding the publication of this article.

## Material and data availability statement

The materials used and datasets generated during and/or analyzed during the current study are available from Professor Bing Hu on reasonable request.

## Appendix Materials & Methods

### Immunostaining

Human oral cavity cancer tissue array that contains normal human gingiva tissues, HnTMA108 was purchased from Creative Bioarray. Slides were heated to 60°C for 20 min before being twice washed in xylenes (Sigma Aldrich 534056) for 10 mins. rehydrated with 100% industrial methylated spirit (IMS) (VWR, 23684.360) for 5 min, before being washed for 2 min in 95% IMS and then 70% IMS. Antigen retrieval was performed by 95°C water bath, slide was submerged in pre-warmed 0.01 M citrate buffer solution (citric acid (Sigma Aldrich, C2404) & 0.05% Tween-20 (Sigma Aldrich, P9416)), pH 8.0 for 20 min. Slide was washed briefly in tap water before washed 3 times in phosphate buffered saline (PBS, Sigma, P4417) containing 0.1% Triton-X100, (Sigma, X100) (PBST) for 5 min per wash. Non-Specific binding was blocked by incubation for 60 min with PBST containing 5% Donkey Serum (Sigma, D9663). Primary antibodies were incubated overnight at 4°C. Slide was washed 3 times in PBST before incubation with secondary antibodies for 2 h at room temperature. Nuclei were counterstained with 2 μg/ml DAPI (Sigma Aldrich, D9542) for 10 min then slide was mounted with Dako fluorescent mounting medium (Dako North America Inc., S3023). IF images were captured using a Leica DMI6000 confocal microscope with a Leica TCS SP8 attachment at a scanning thickness of 1 μm per section. The microscope is running LAS X (Leica, Version 3.7.3.23245) software from Leica. Images for comparison were taken using the same settings and post imaging processing was conducted using Adobe Photoshop (Adobe, Version 24.0.1).

For frozen section, the slides were air-dried for 1 h then fixed with 4% paraformaldehyde (PFA; Sigma in PBS) for 30 min, then washed twice with PBST 5 min each. Primary antibodies were incubated overnight at 4°C. Slides were washed three times in PBST before incubation with secondary antibodies for 2 h at room temperature. Nuclei were counterstained with 2 μg/ml DAPI for 10 min.

Images were captured using a Leica DMI6000 confocal microscope with a Leica TCS SP8 attachment. The microscope is running LAS X (Leica, Version 3.7.3.23245) software. Images for comparison were taken using the same settings and post imaging processing was conducted using Adobe Photoshop (Adobe, Version 24.0.1). Images of 3D equivalents were visualized by Imaris software (Bitplane, Version 9.0.2).

### BrdU incorporation assay

5*10^3 of HPLFs were seeded into a black 96 well plate (GBO, 655090). After the spike protein treatment or lentiviral infection, BrdU cell prolife labelling reagent (Amersham-GE, RPN201, 1:1000) was diluted in complete cell culture medium then added into dishes for 2-3 h. Cells were washed 3 more times with PBS for 2 min each, then fixed in 4% PFA for 30 min followed by washing twice in PBS. Cells were treated with 100 μl 2N hydrochloric acid (HCl) (Sigma, H1758-100ml) for 30 min before they were stained with anti-BrdU antibody (Abcam, ab6326, 1:500). The antibody staining procedures were exactly as above except that the blocking buffer was prepared with 2.5% bovine serum albumin (BSA) (Sigma. A2153) in PBST. Images were processed in Image J software (National Institutes of Health, Bethesda, MD, USA, Version 1.53k). BrdU-positive cells and total cell number were quantified using ‘Analyze Particles’ function.

### Terminal deoxynucleotidyl transferase dUTP nick end labelling (TUNEL) assay

5*10^3 of HPLFs were seeded into black 96 well plate. After the spike protein treatment or lentiviral infection, cells were washed briefly for 5 sec in PBS twice, then fixed in 4% PFA for 30 min and washed twice in PBS 5 min each. Cells were permeabilized by PBST for 2 min on ice then incubated in 50 μl TUNEL mixture (In Situ Cell Death Detection Kit, Fluorescein, version 17, Roche) for 2 h at 37°C in humid atmosphere. After the washing and mounting, images were taken by Leica IM8 conducted using Adobe Photoshop.

### Senescence assay

5*10^4 of HPLFs were seeded in 24 well plate. After the spike protein treatment or lentiviral infection, cells were washed briefly with PBS, the fixative solution from the senescence kit (Senescence b-galactosidase kit, Cell Signalling, 9860S) was added to each well for 15 min at room temperature. Cells were rinsed 2 times with PBS then β-Galactosidase Staining Solution, pH 6.0, was added into each well, and at the meantime pH 4.0 was added into spare wells as positive control. The plate was incubated at 37°C at least overnight in a dry incubator without CO2. Cells were washed twice with PBS then stained with nuclear fast red solution (Sigma, N8002) for 20 min then washed with distilled H2O. All wells were mounted with 70% glycerol (Sigma, G2025) and images were taken with Leica IM8 conducted using Adobe Photoshop.

### Proteomic analysis

HPLFs were seeded in 6cm dishes and performed viral infection and protein treatment for 6 h and 48 h. To collect protein after each time points, culture media was removed from the culture dishes and cells were washed in pre-cooled HBSS twice. HBSS was removed and replaced with ice-cold RIPA buffer (ThermoFisher, 89901) supplemented with Halt™ Protease and Phosphatase Inhibitor Cocktail (ThermoFisher , 78440) at 1:100. The cells were detached from the dish using a cell scraper and then collected into an Eppendorf tube on ice then incubated for 30 min with frequent agitation for efficient cell lysis and solubilisation of proteins. Tubes were spun down at 15,000 rpm for 15 min at 4°C so the supernatant containing the protein could be collected and stored at -80°C until ready to load onto a gel. 15 µg protein samples were run on a Nupage 4-12% Bis-Tris protein gel at 200V for 45 min. Gel was rinsed with water before fixing in 40% ethanol and 10% acetic acid for 15 min with gentle agitation. After washing the gel twice in water, the gel was stained overnight in QC colloidal Coomassie Blue G-250 (Biorad, 161-0803). The gel was destained for 1 h with changes of water every 15 min until protein bands were visible. Every sample lane was cut into 4 fractions and each fraction further cut into 1-2 mm cubes for equilibration. In-gel digestion, sample cleanup and mass spectrometric analysis was carried out as described previously (Dunn, J et al, 2018). All samples were stored at -20°C or analyze directly using mass spectrometry.

## Appendix Figures

**Appendix Figure 1.**
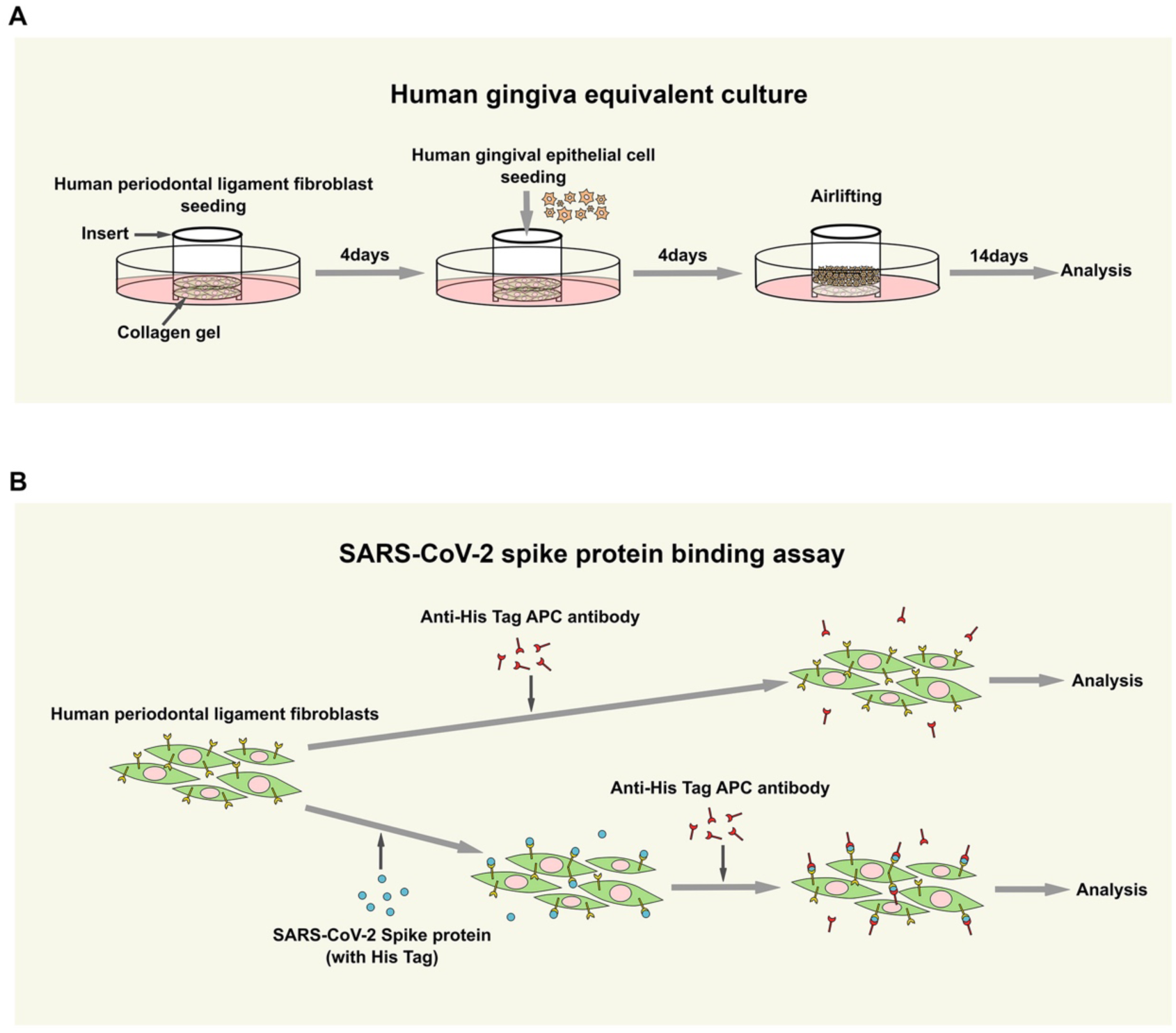
Illustration of human gingiva equivalent culture and spike protein binding assay. A. For human gingiva equivalent culture, HPLF were seeded into collagen gel supported by a cell culture insert cylinder. 4 days later human gingival epithelial cells were seeded on top of the culture. After a further 4 days, the culture were lifted to air-liquid interface to allow stratification for 14 days before further analysis. B. To test the binding of spike protein to HPLF cells, the cells were treated with His Tag conjugated spike protein first then traced using anti-His Tag APC conjugated antibodies. For control, the spike proteins were omitted.

**Appendix Figure 2.**
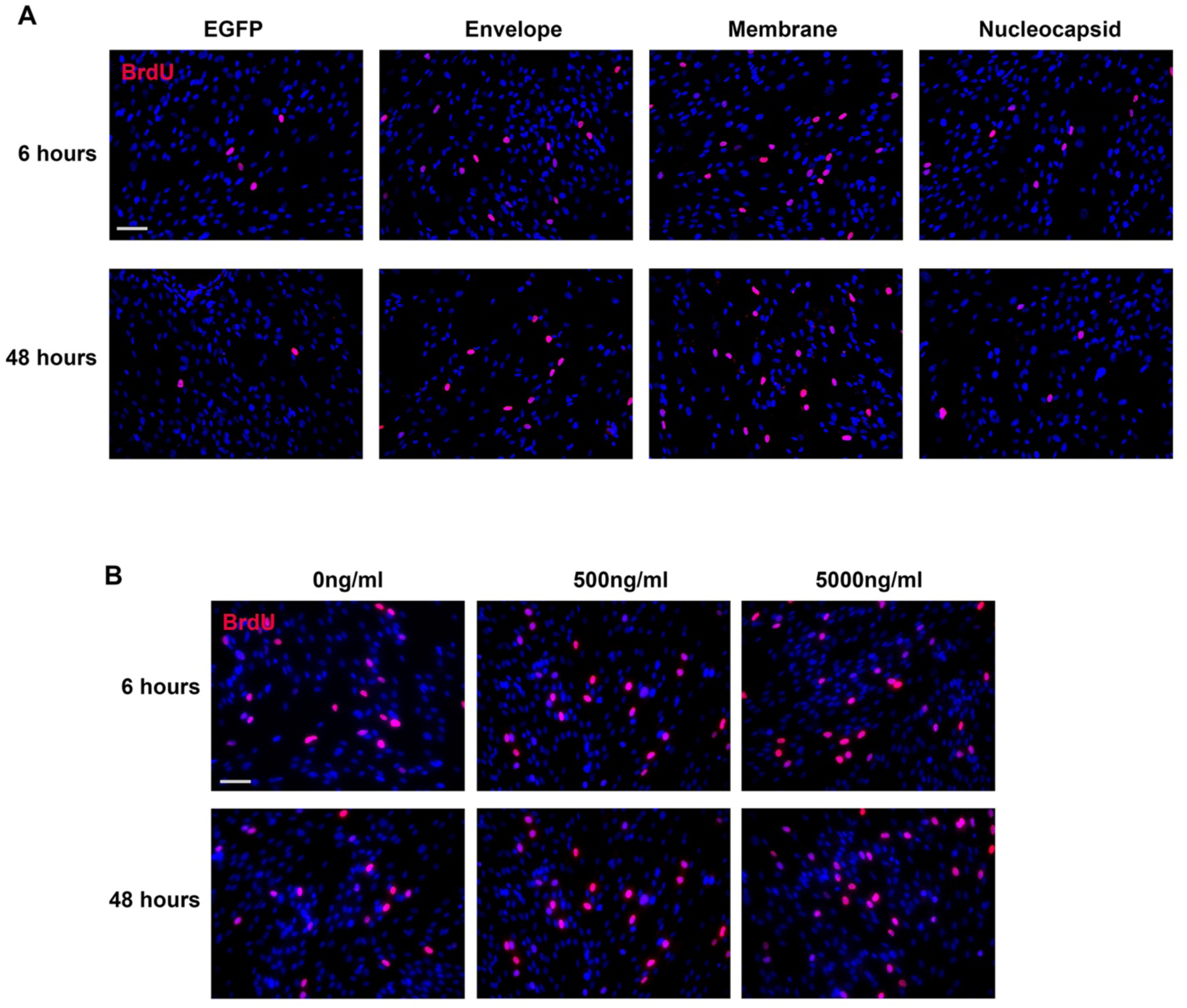
SARS-CoV-2 envelope and membrane protein could induce PDL fibroblast proliferation. Representative field images of BrdU incorporation analysis in the indicated conditions. Quantitative analysis can be found from Figure 2 A-D. Bars: 100um

**Appendix Figure 3.**
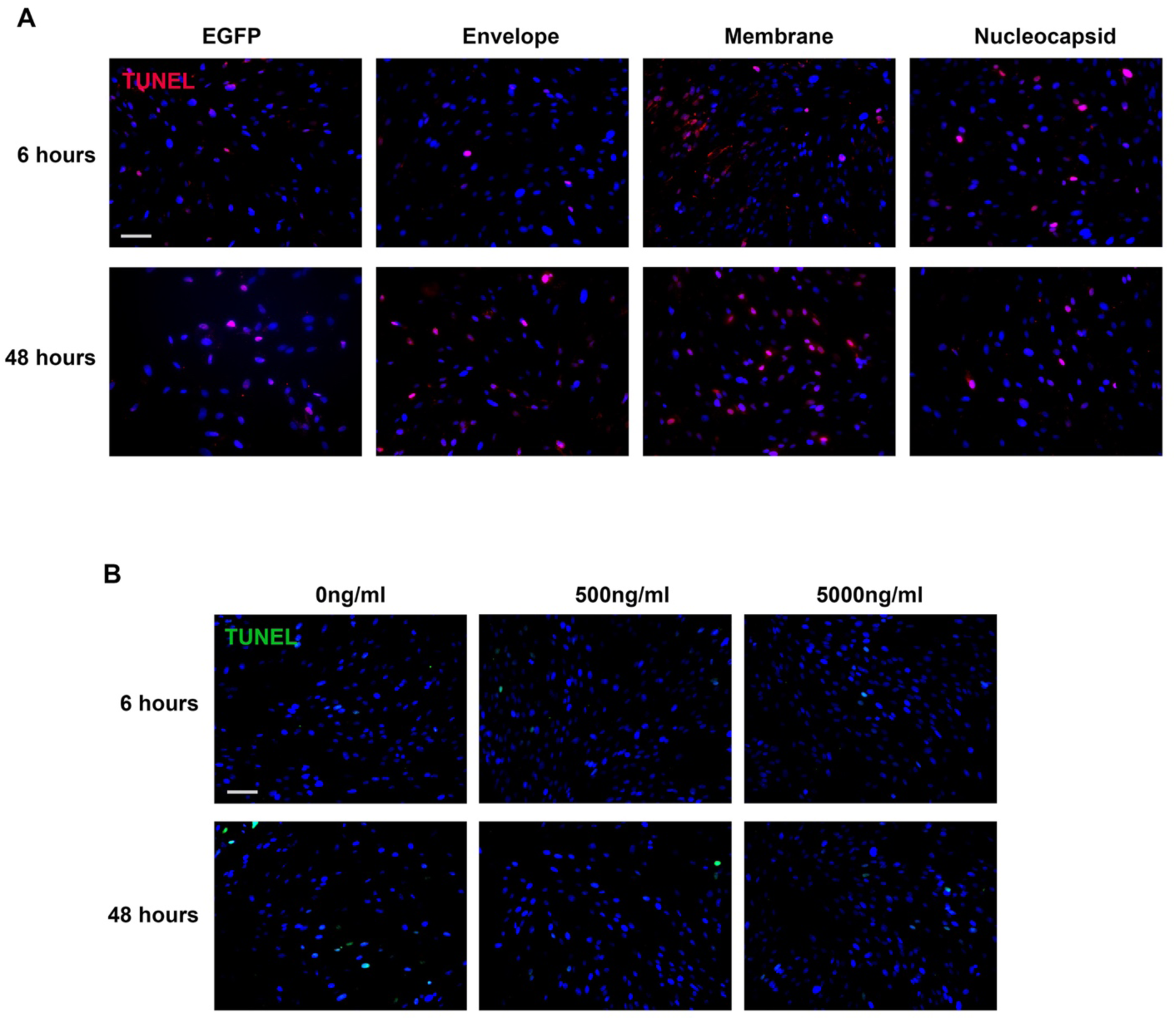
SARS-CoV-2 envelope and membrane protein could induce PDL fibroblast apoptosis. Representative field images of TUNEL analysis in the indicated conditions. Quantitative analysis can be found from Figure 2 E-H. Bars: 100um

**Appendix Figure 4.**
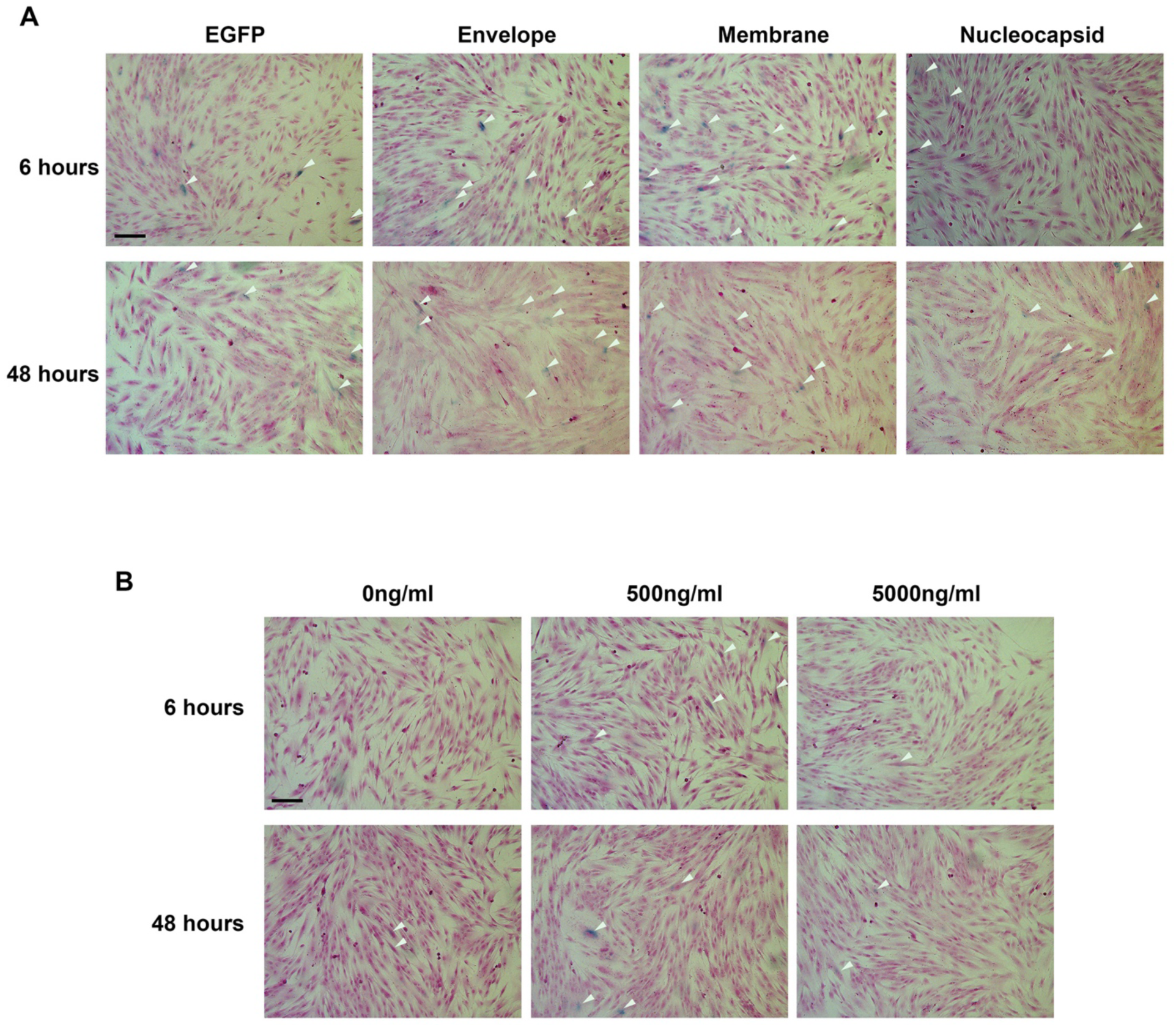
SARS-CoV-2 envelope and membrane protein could induce PDL fibroblast senescence. Representative field images of senescence analysis in the indicated conditions. The cells were countered stained with nuclear fast red. Note increased blue staining (marking senescent cells) in the envelope and membrane groups. Quantitative analysis can be found from Figure 2 I-L. White arrows indicate representative positive cells. Bars: 100um

**Appendix Figure 5.**
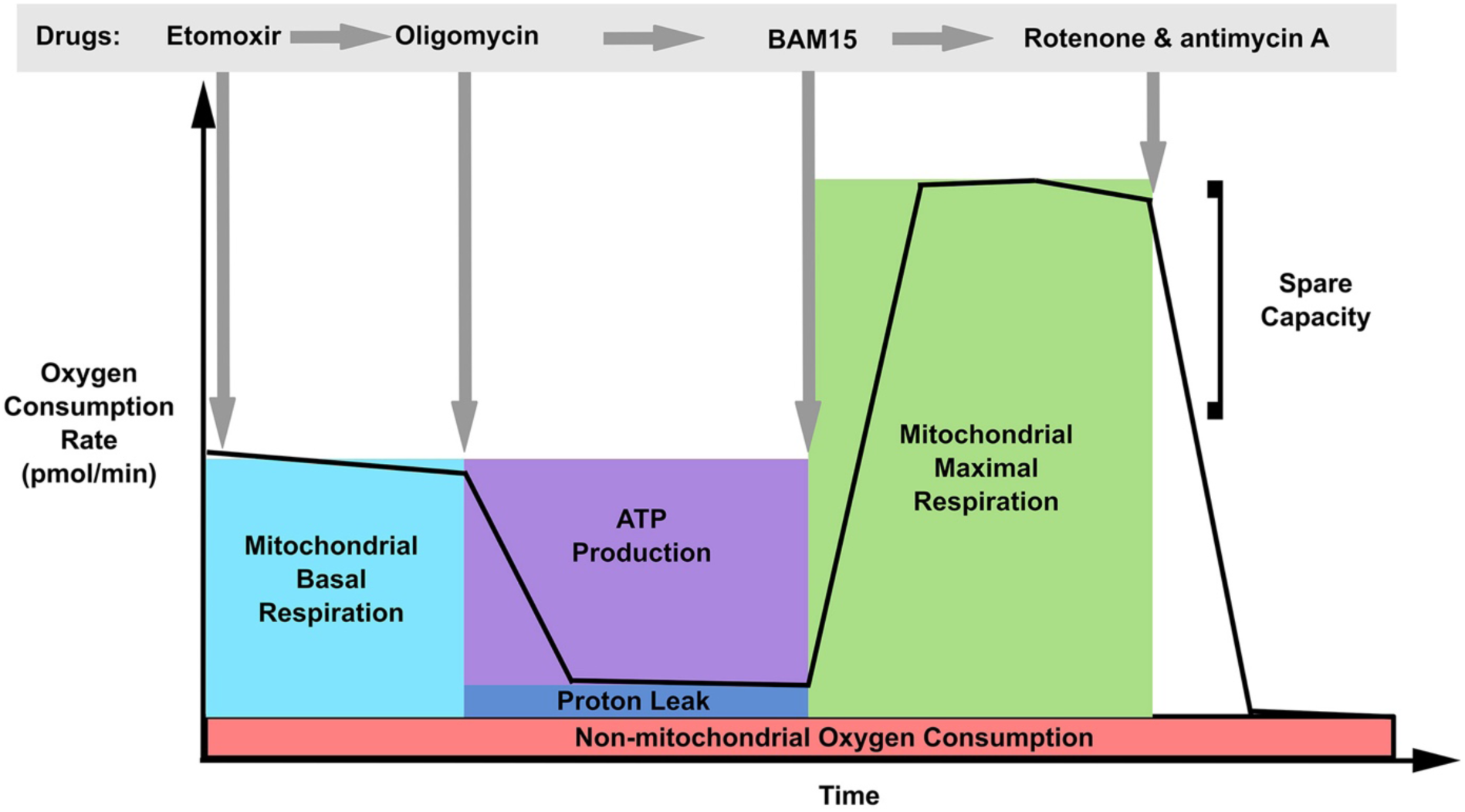
Illustration of Seahorse mito stress test, for which drugs were added sequentially into the cell culture and what have been measured by the machine.

**Appendix Figure 6.**
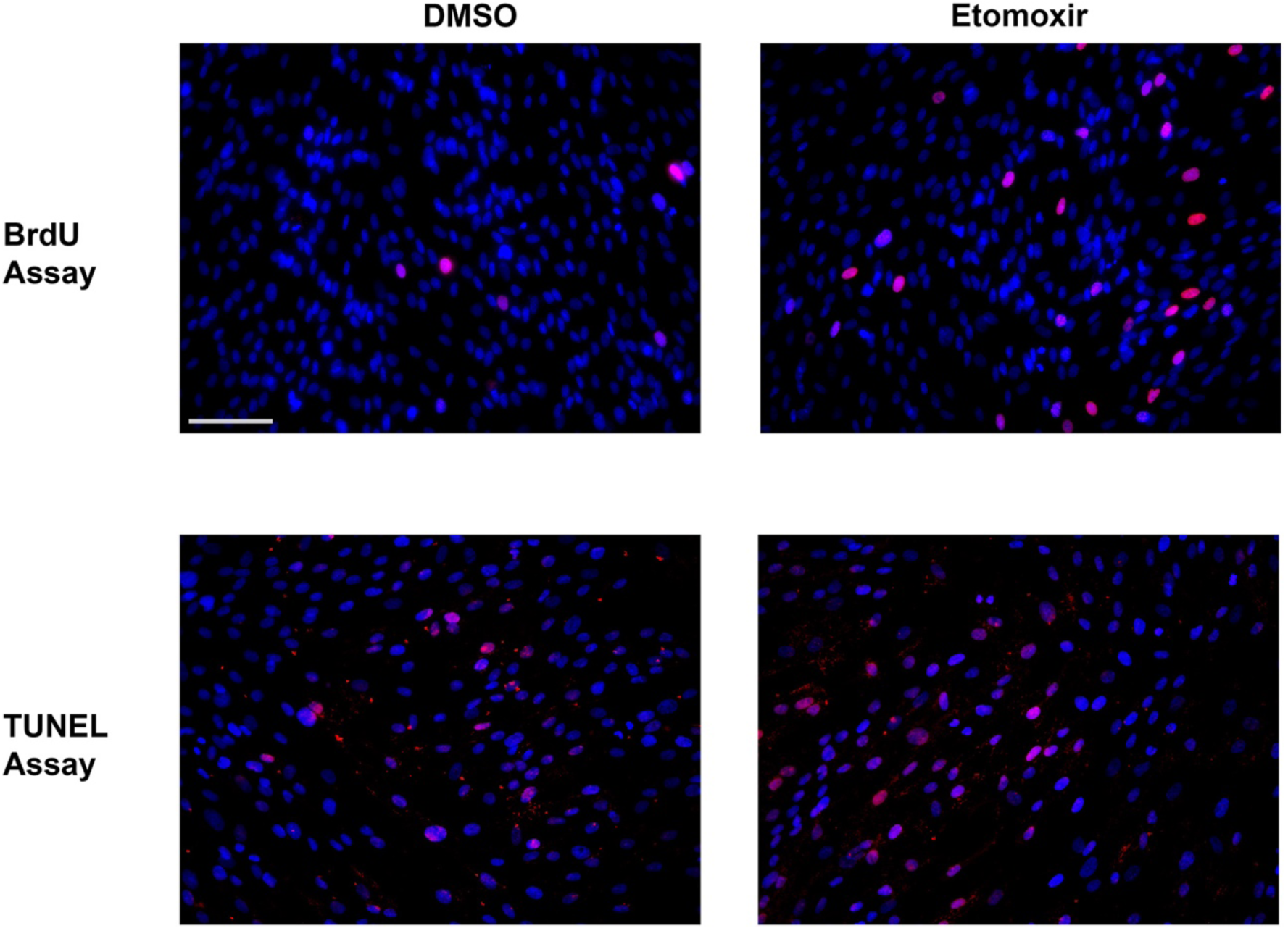
Mitochondrial fatty acid β-oxidation inhibition mirrored fibrotic degeneration phenotypes in PDL fibroblasts. Representative field images of BrdU incorporation and TUNEL analysis in the indicated conditions. Quantitative analysis can be found from Figure 5 B and C.

**Appendix Table 1.**
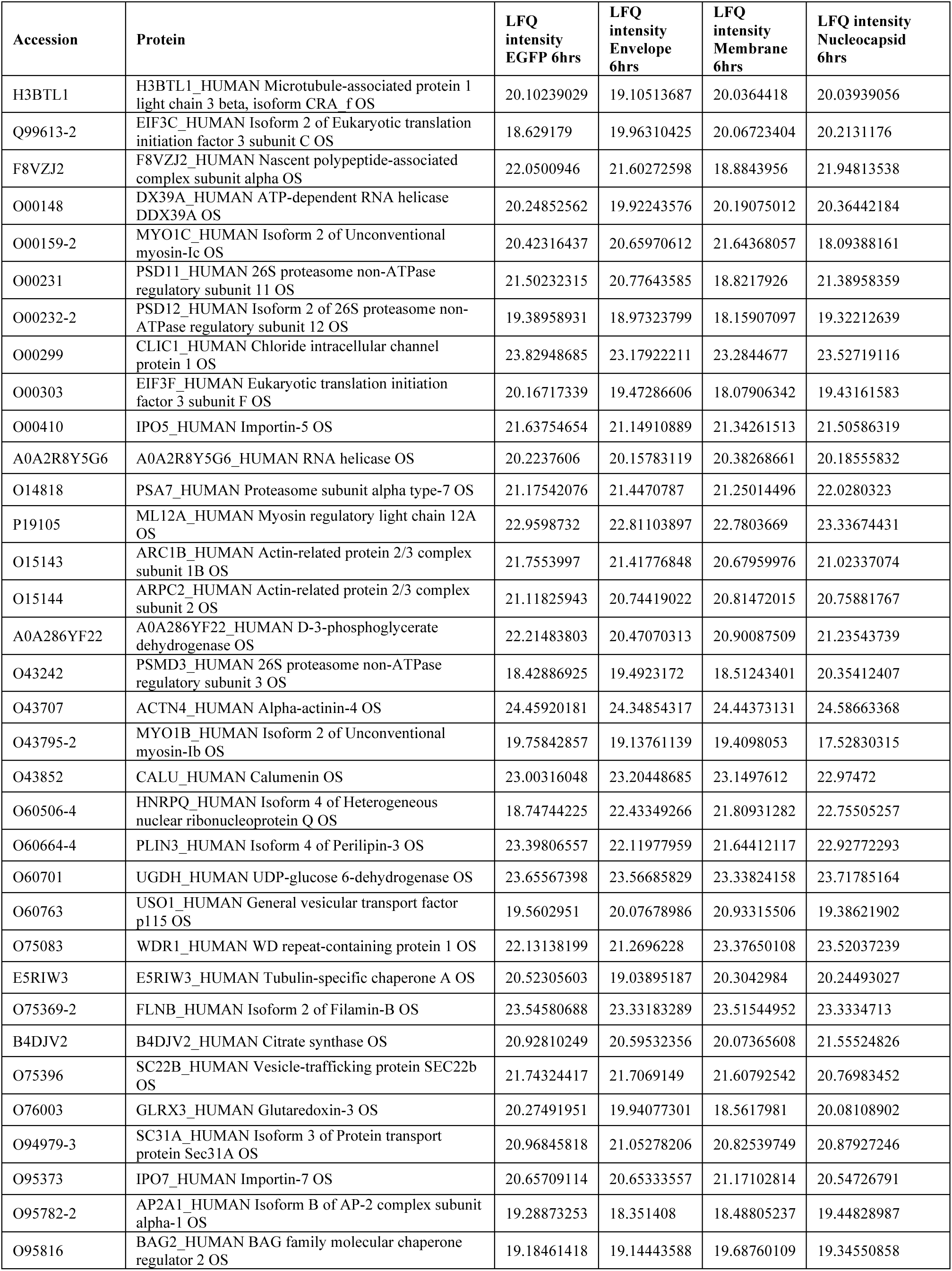
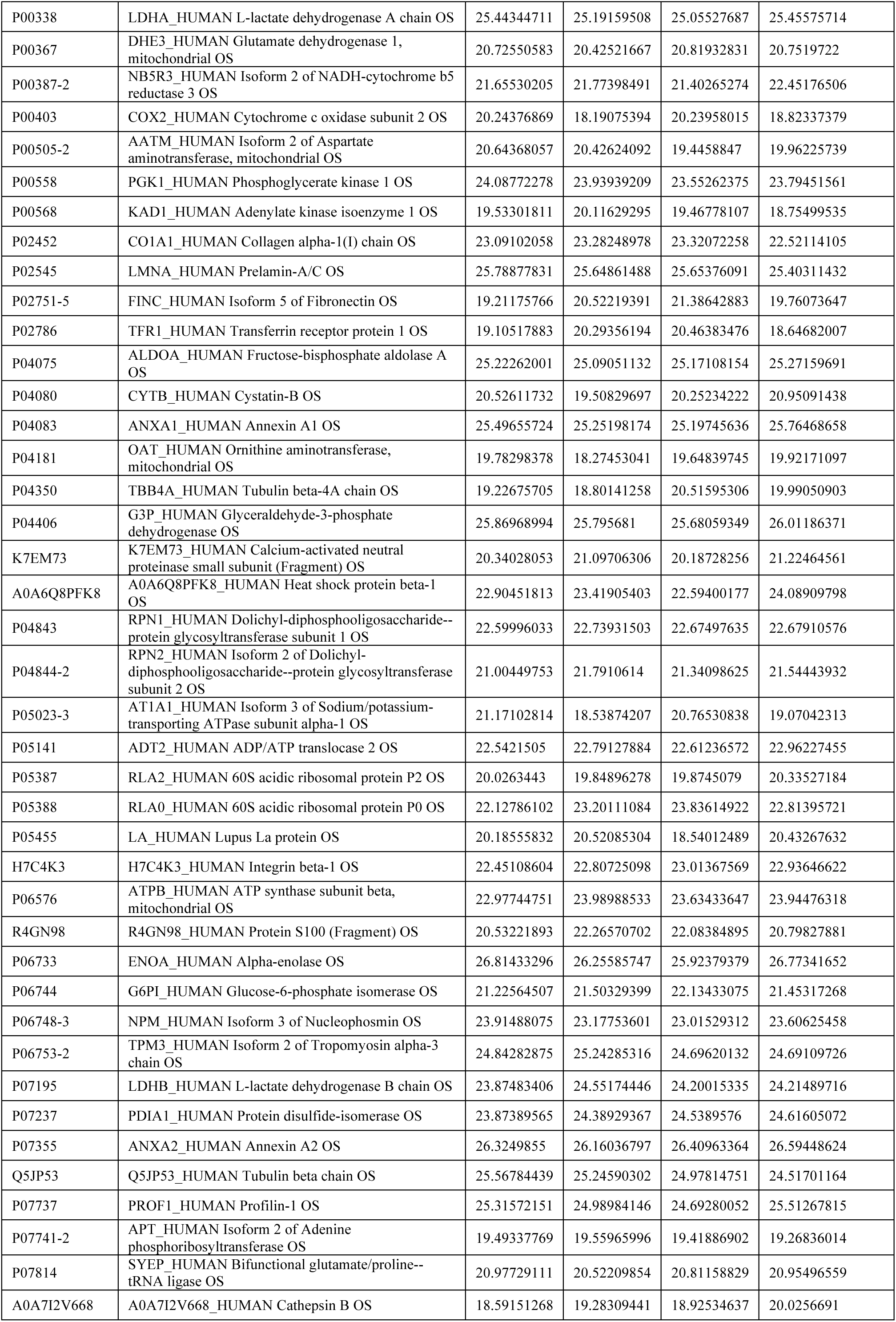
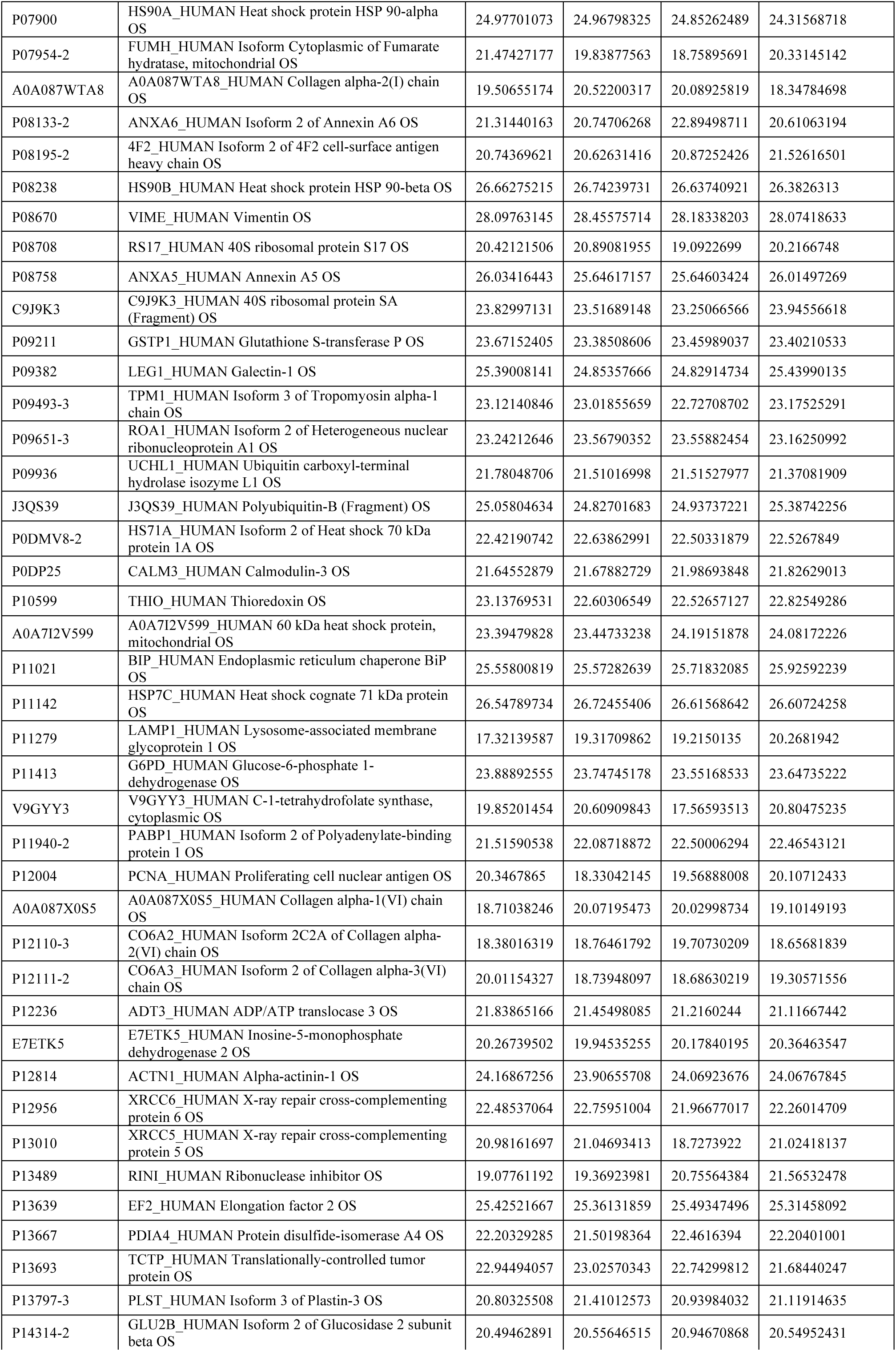
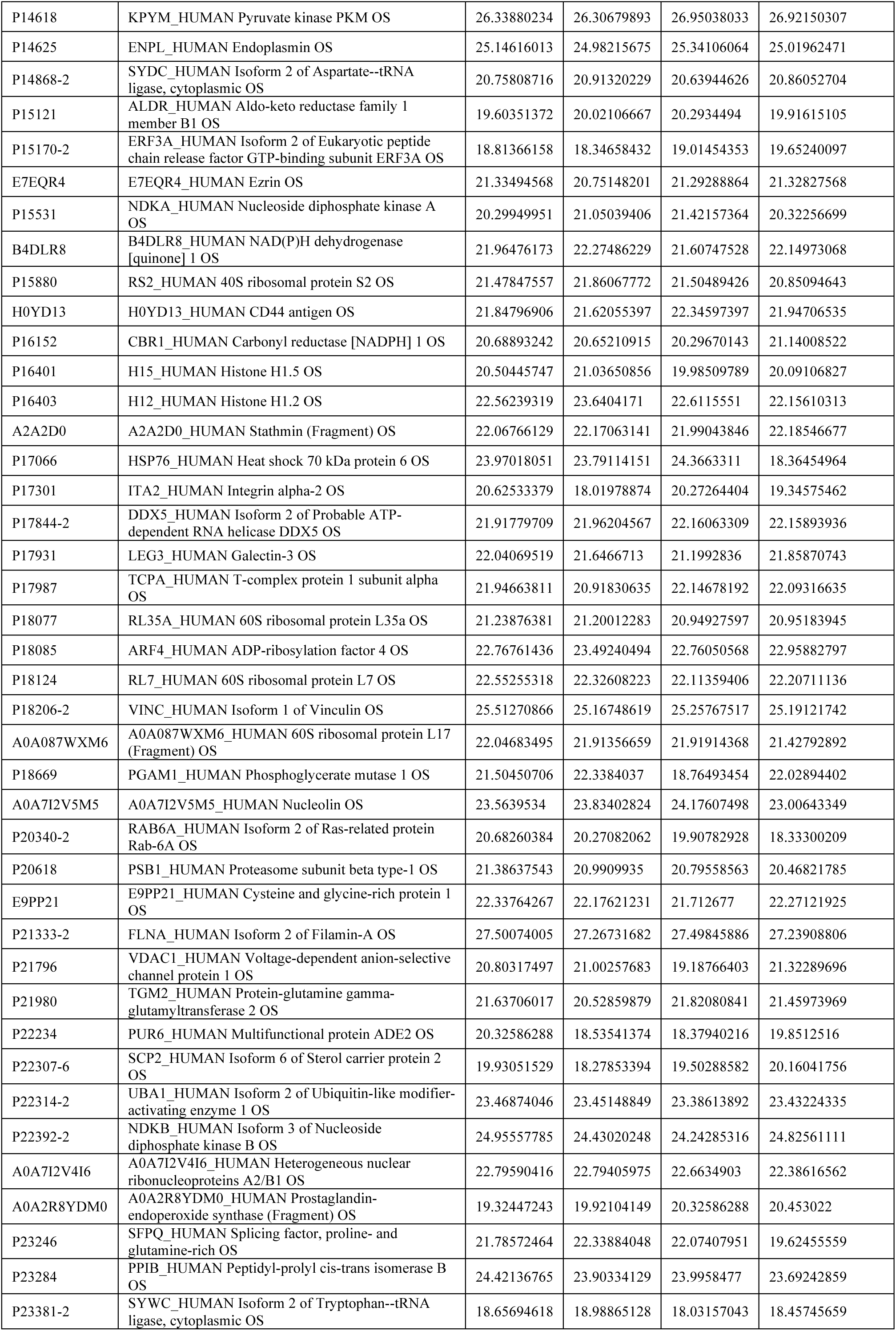
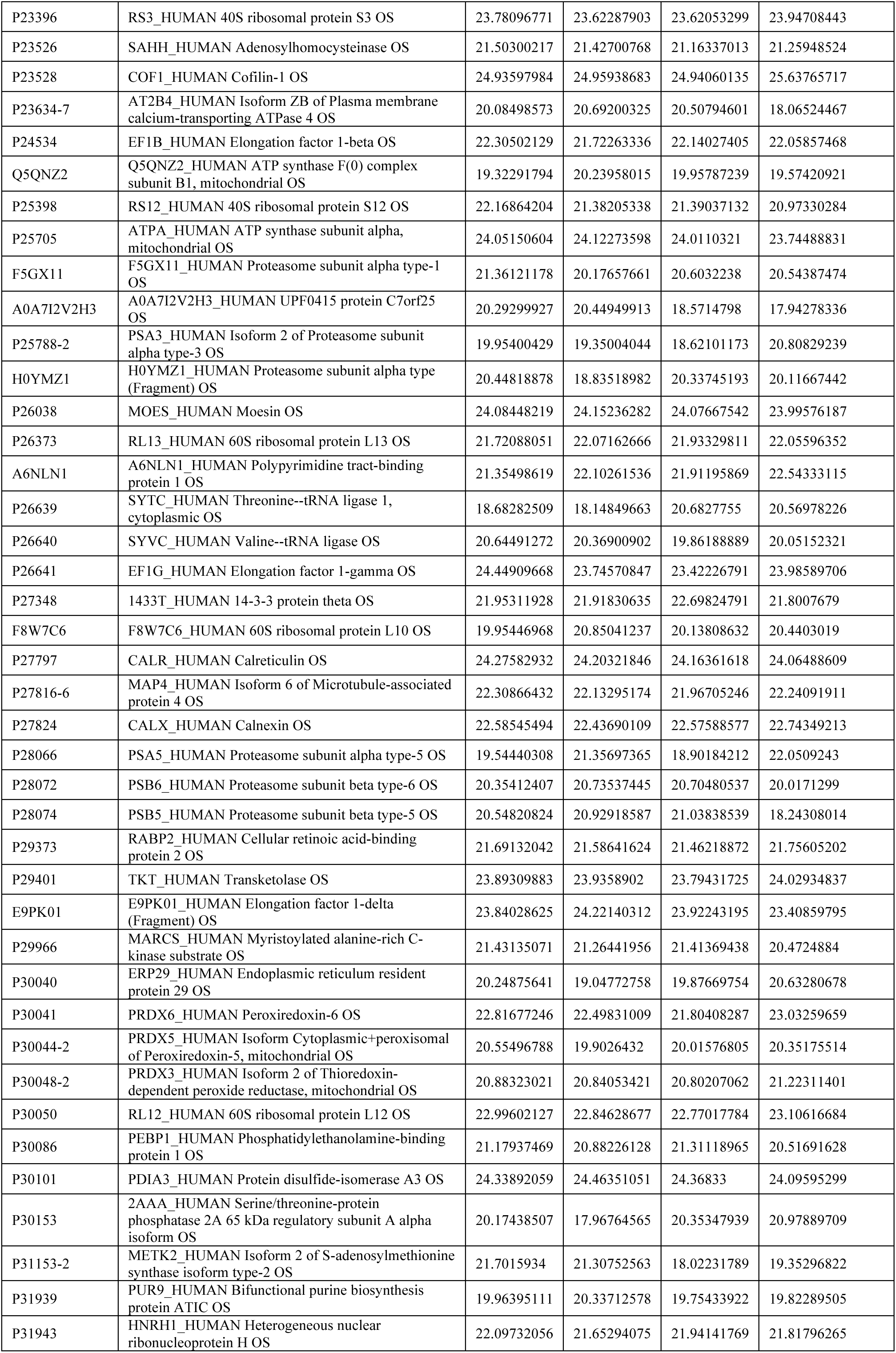
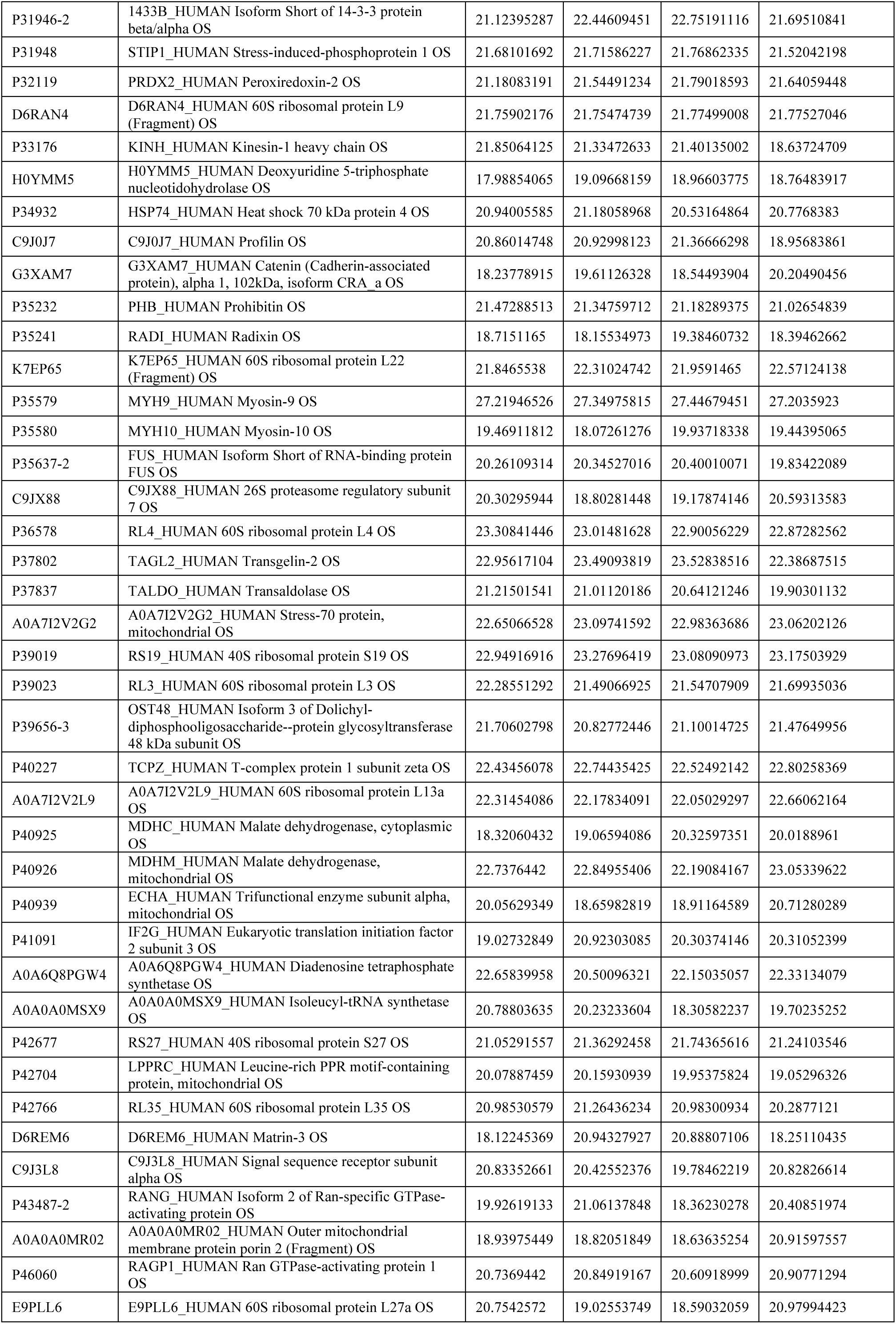
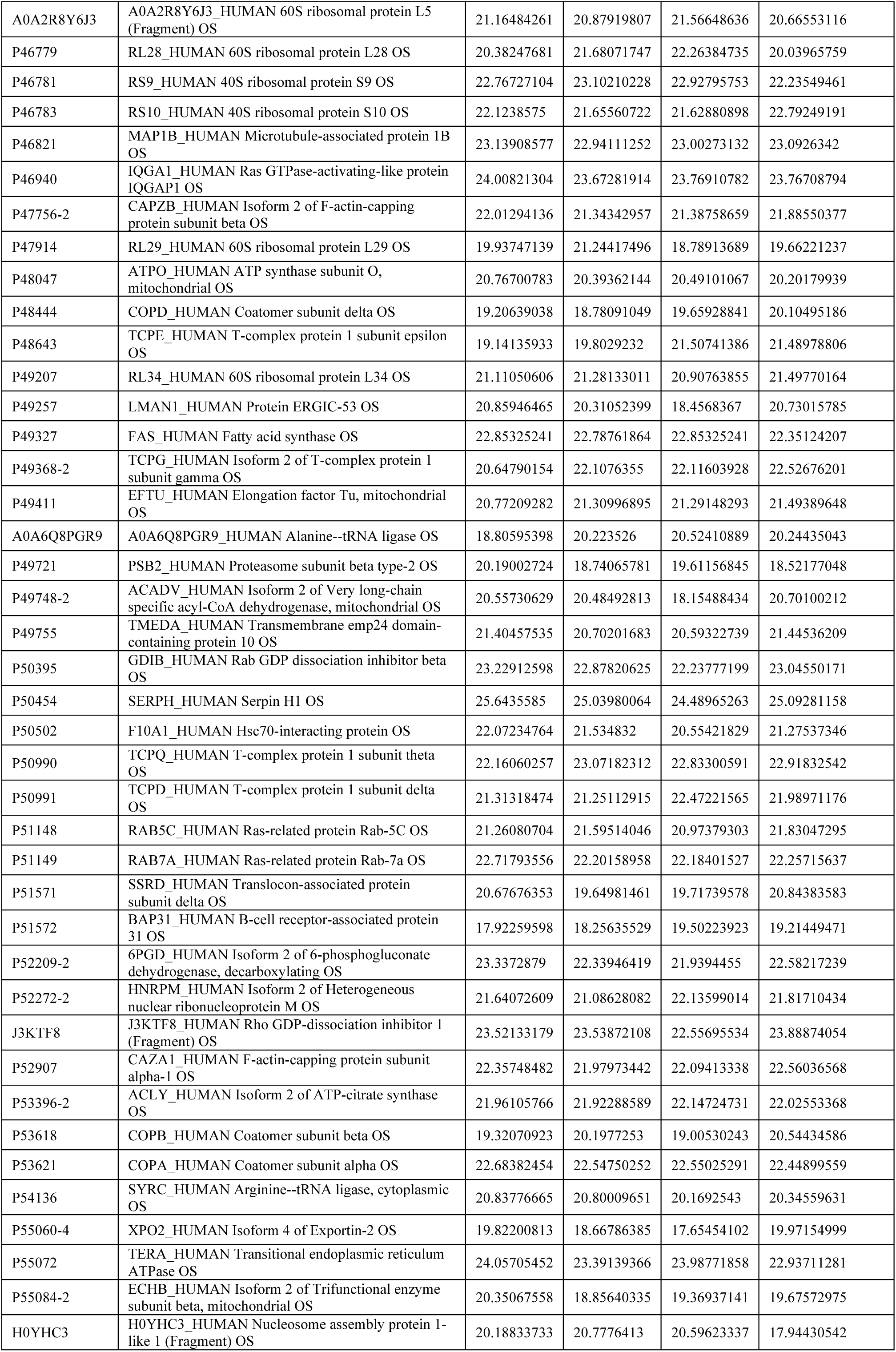
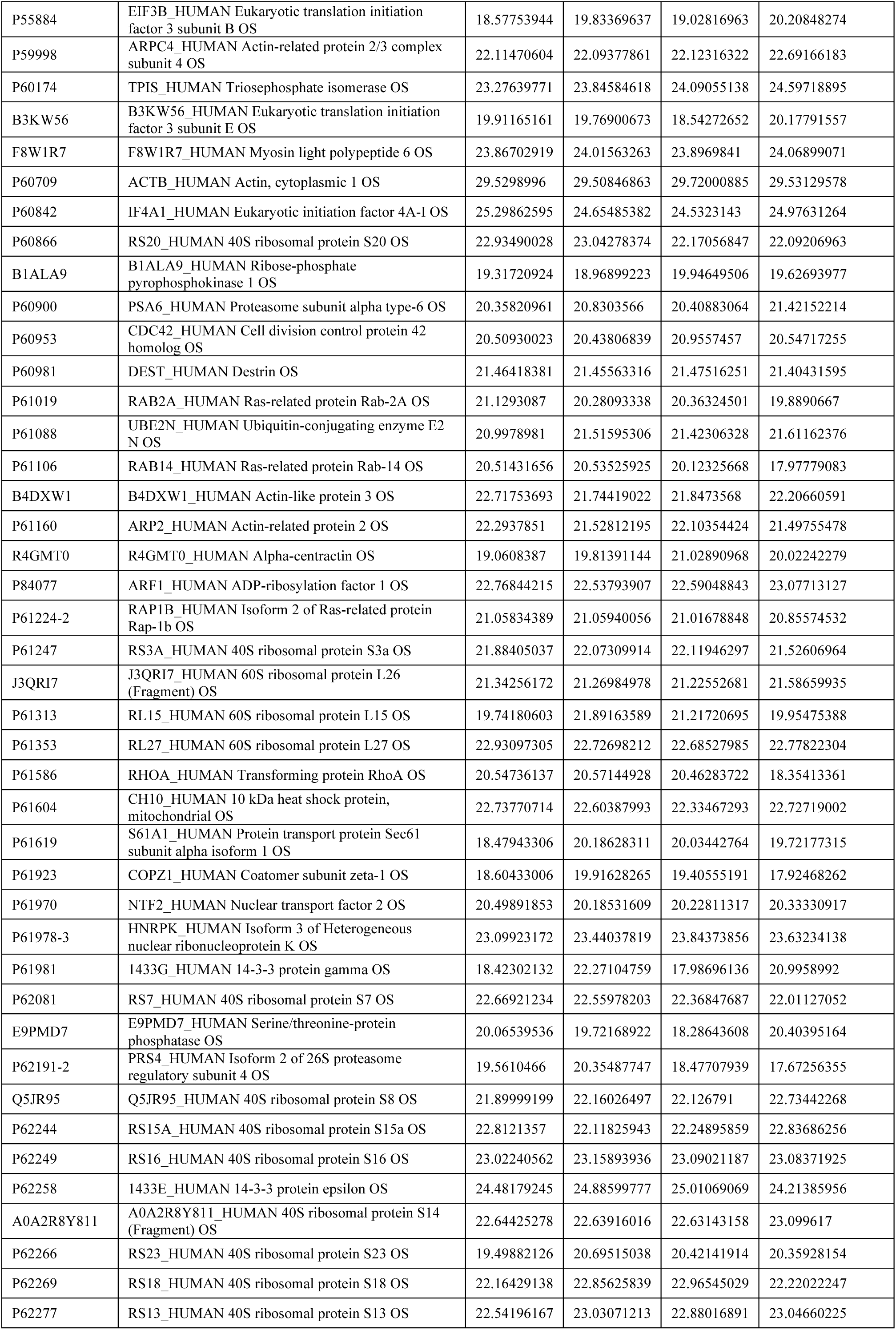
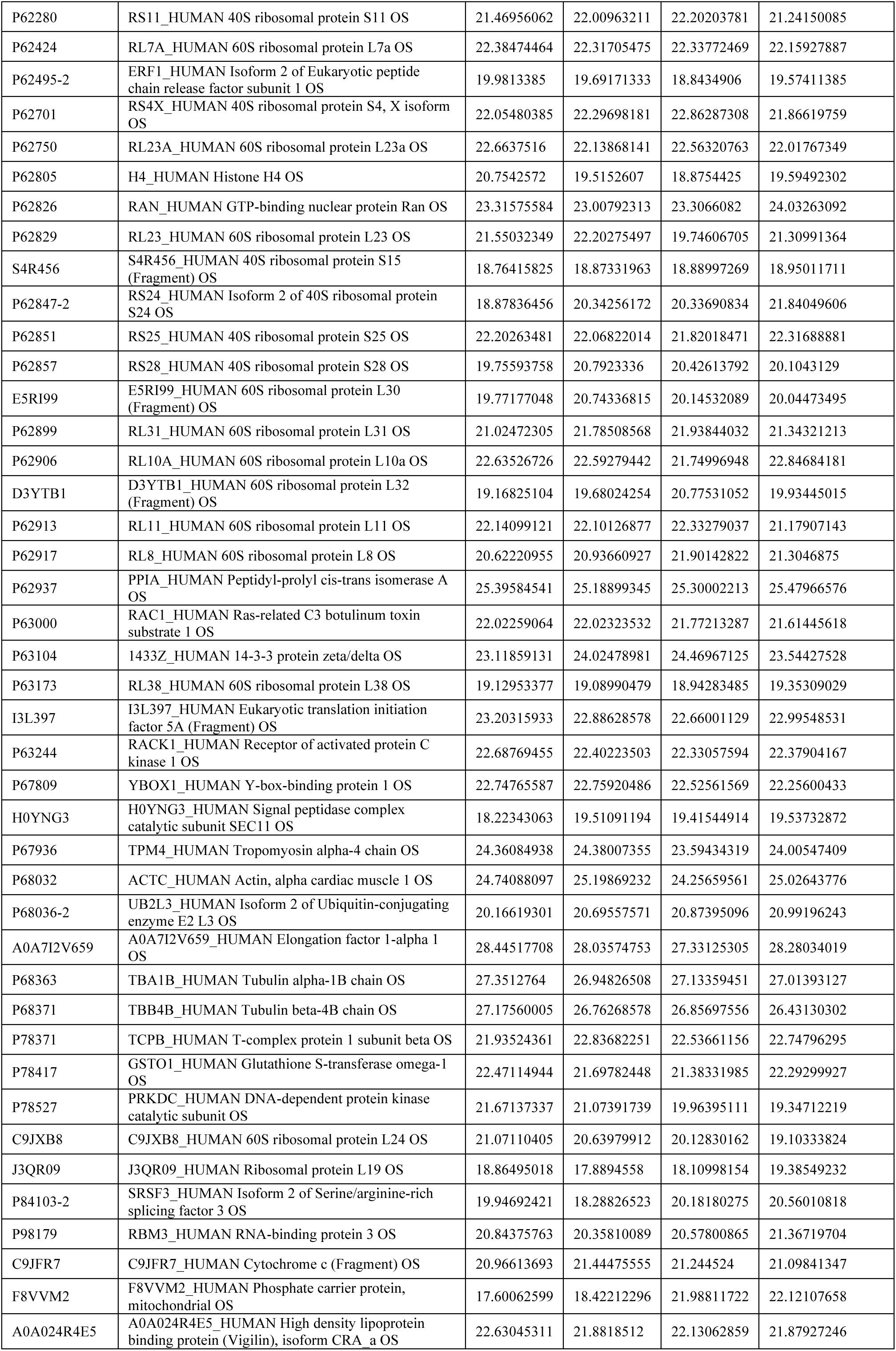
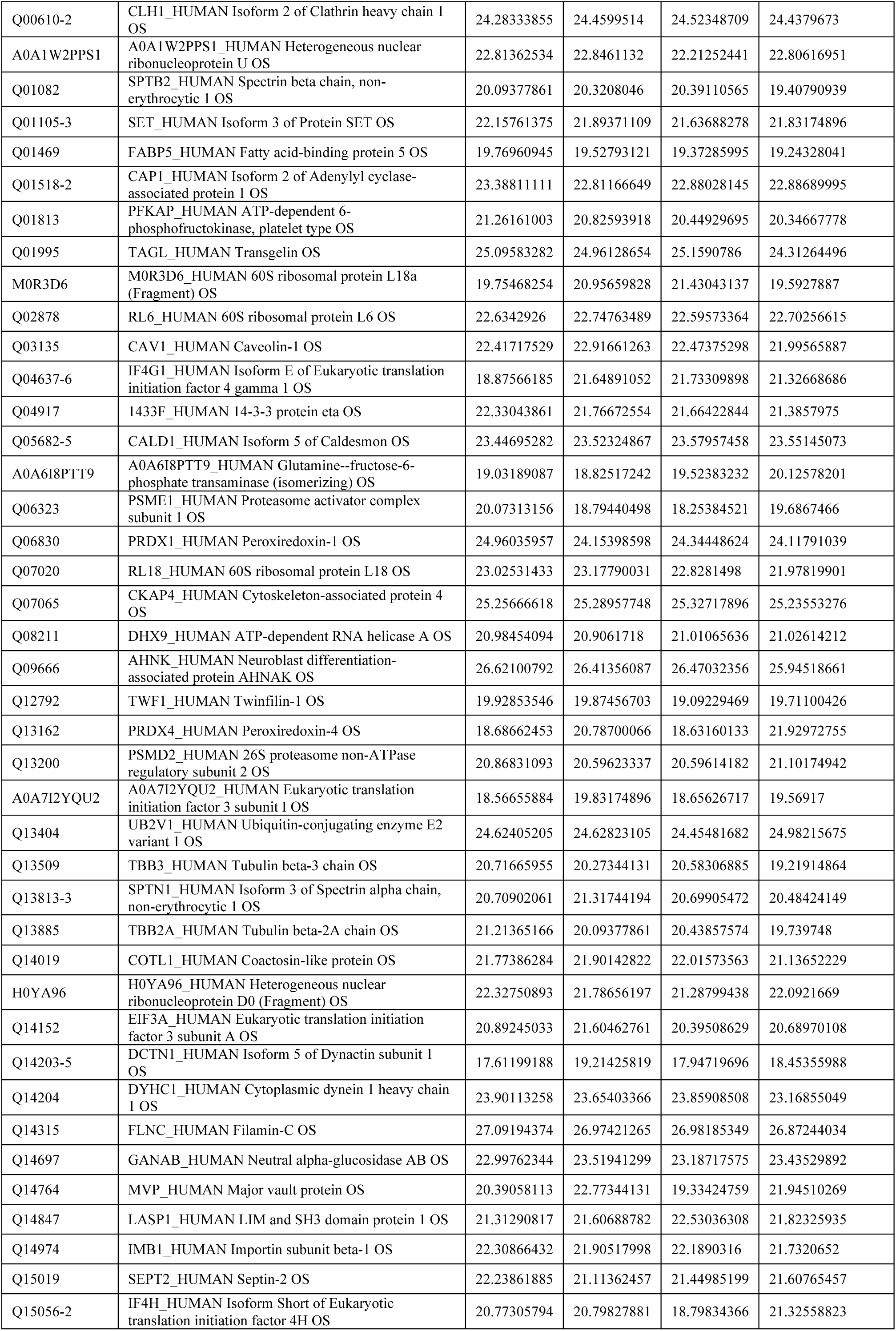
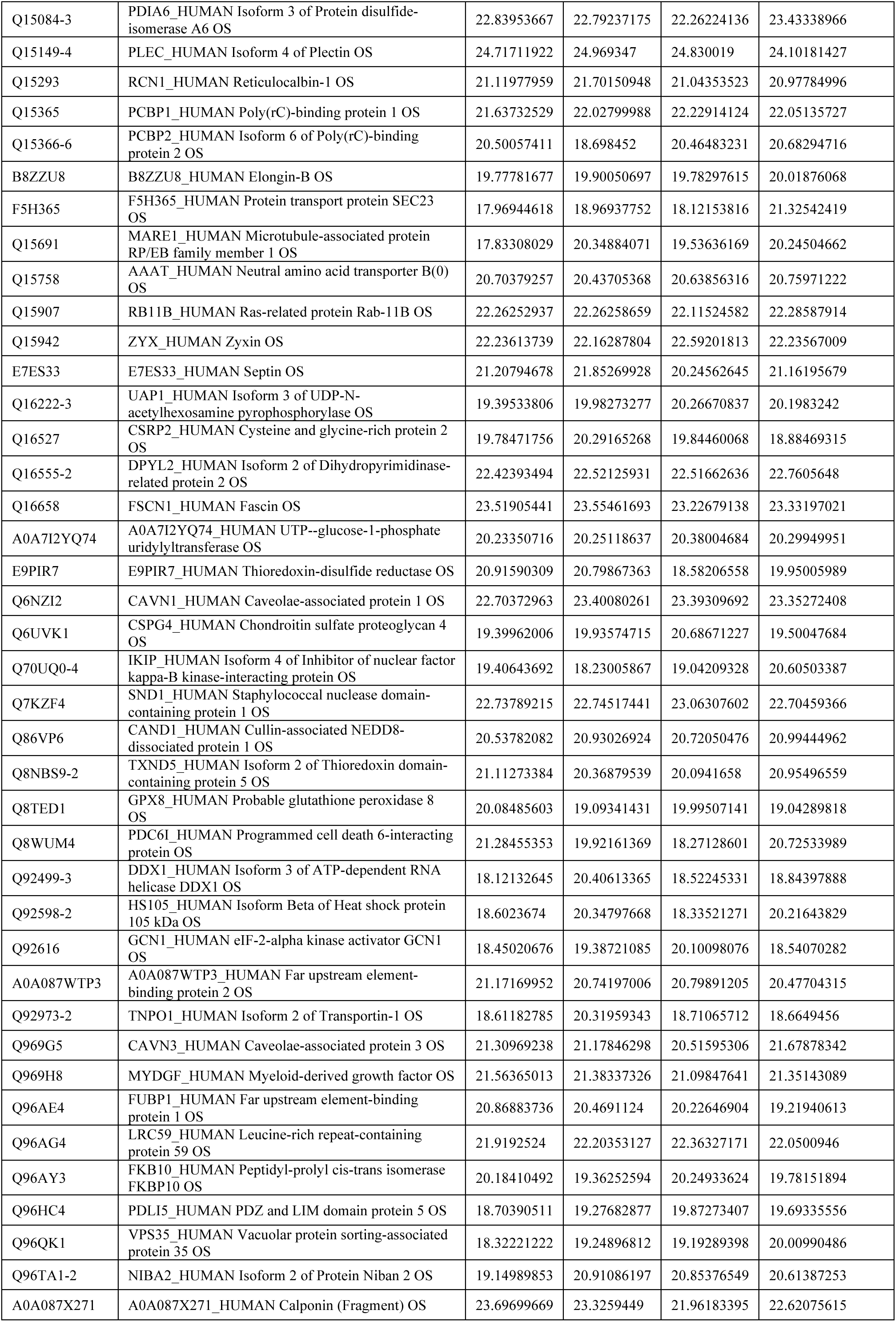
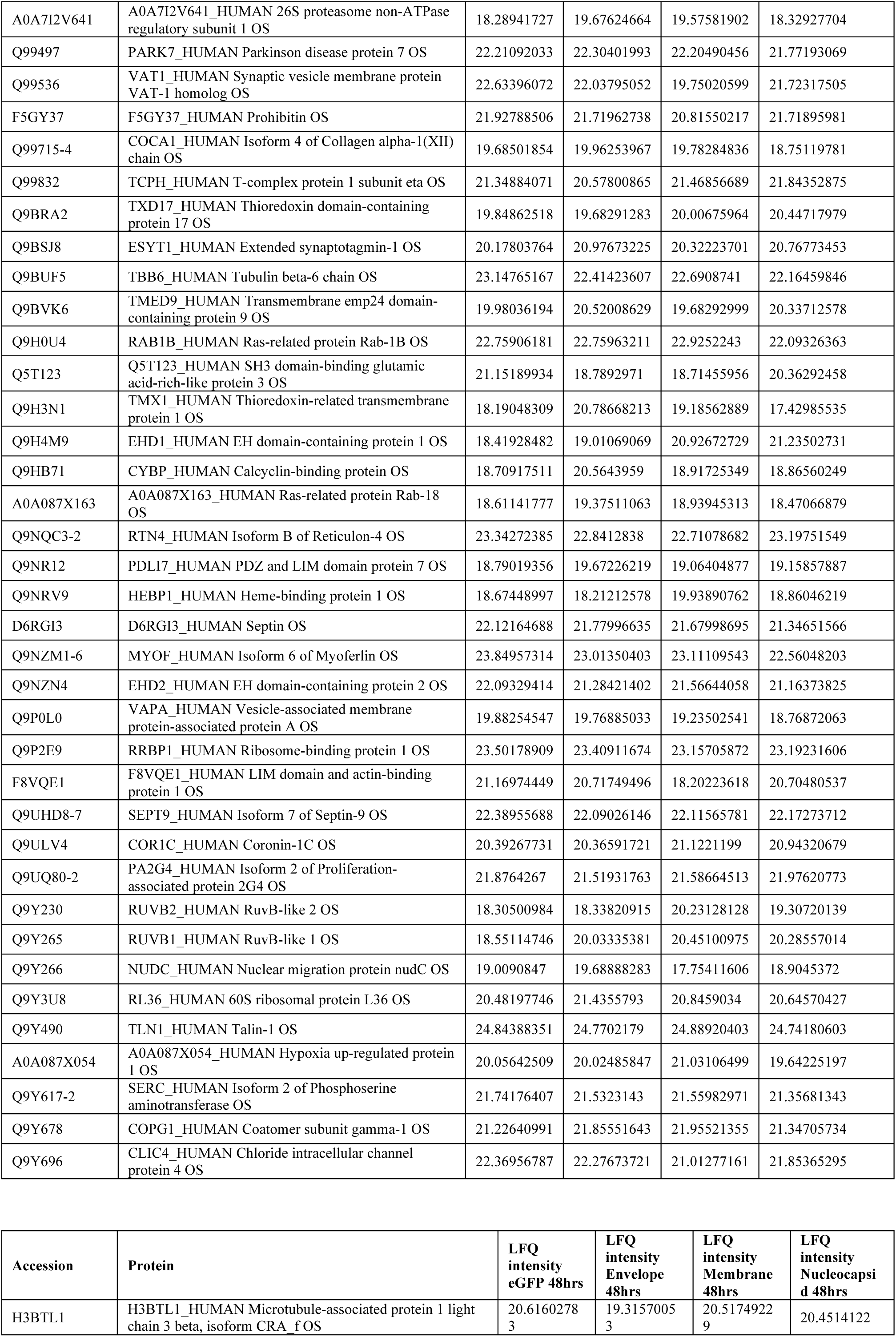
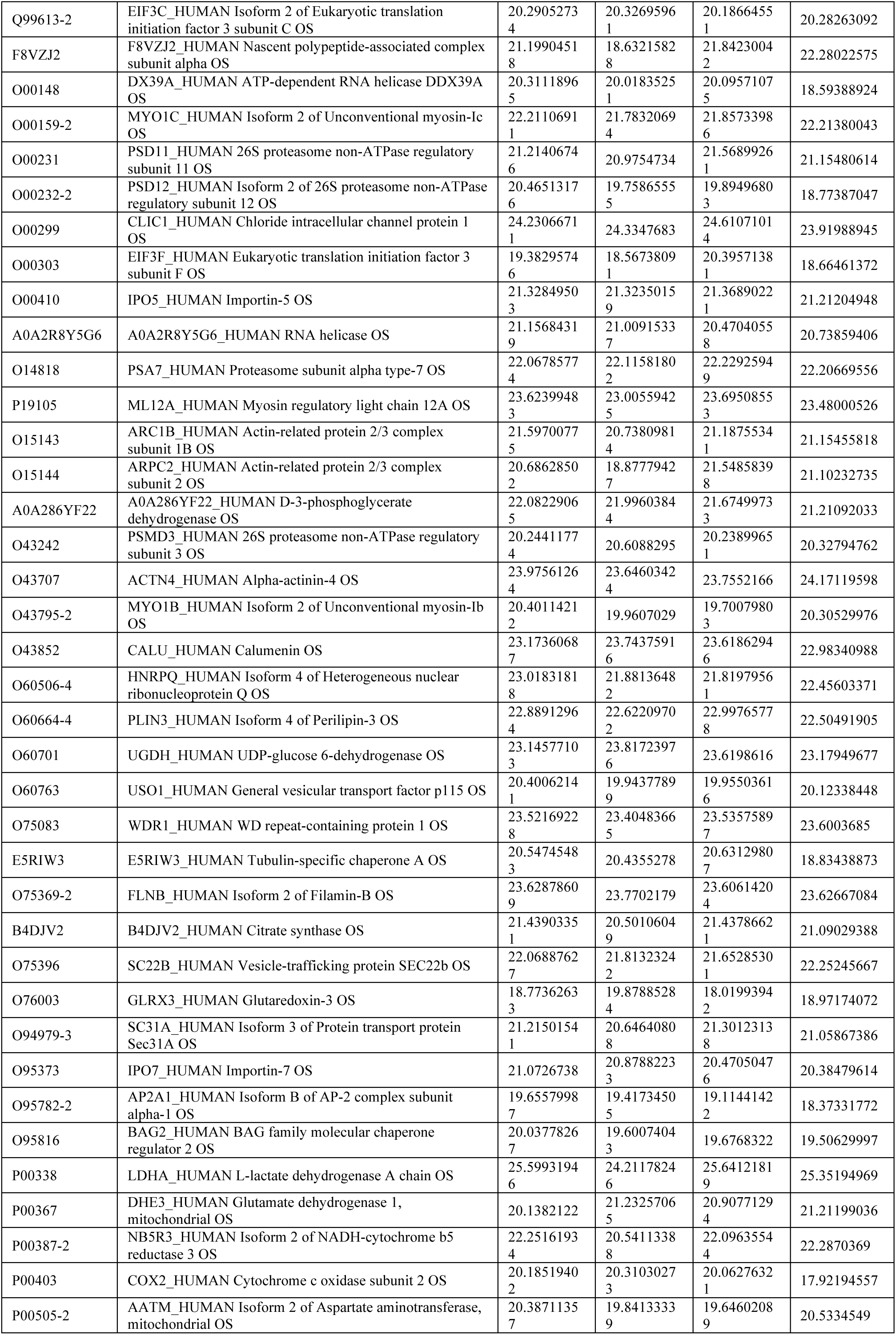
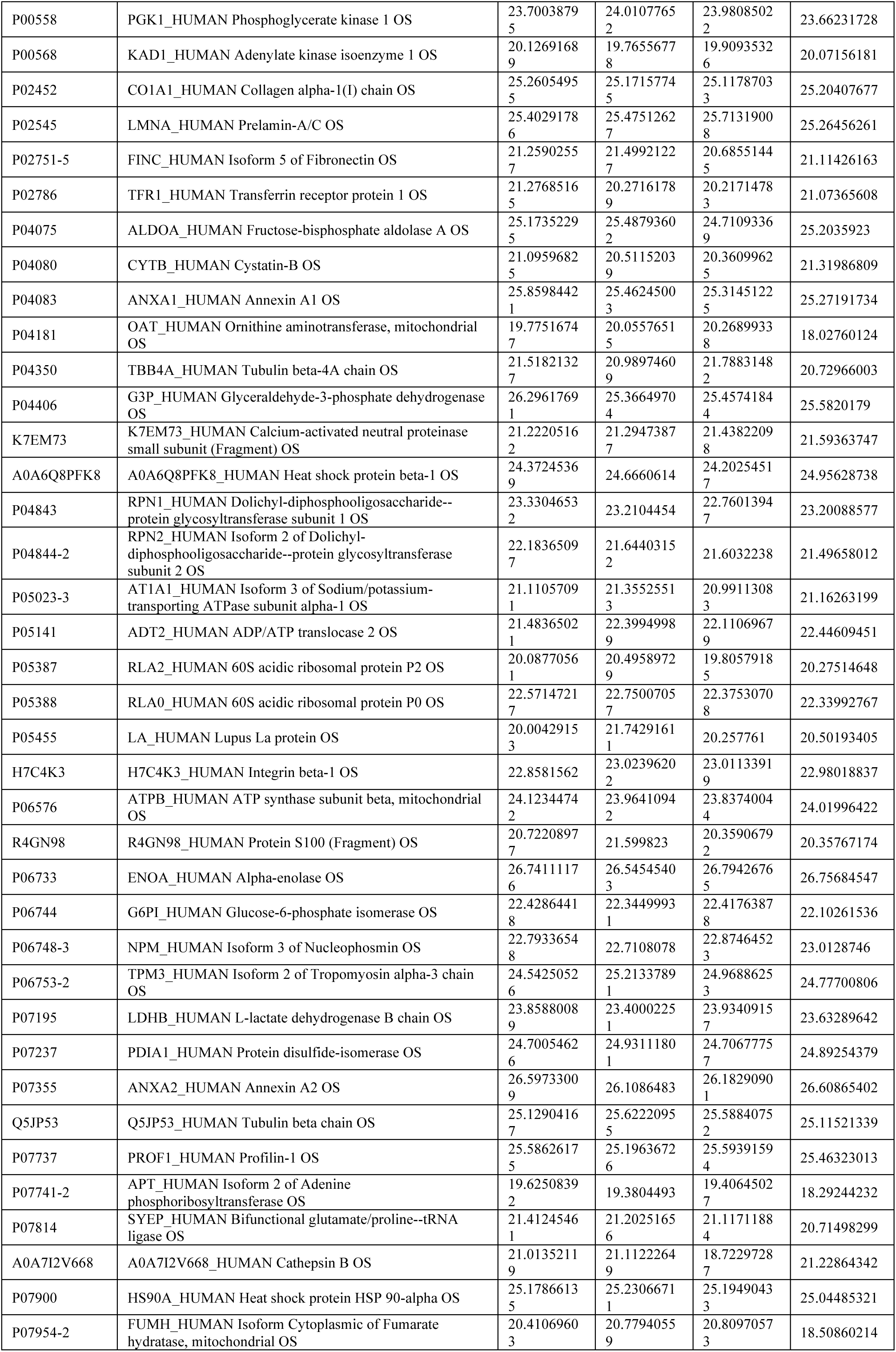
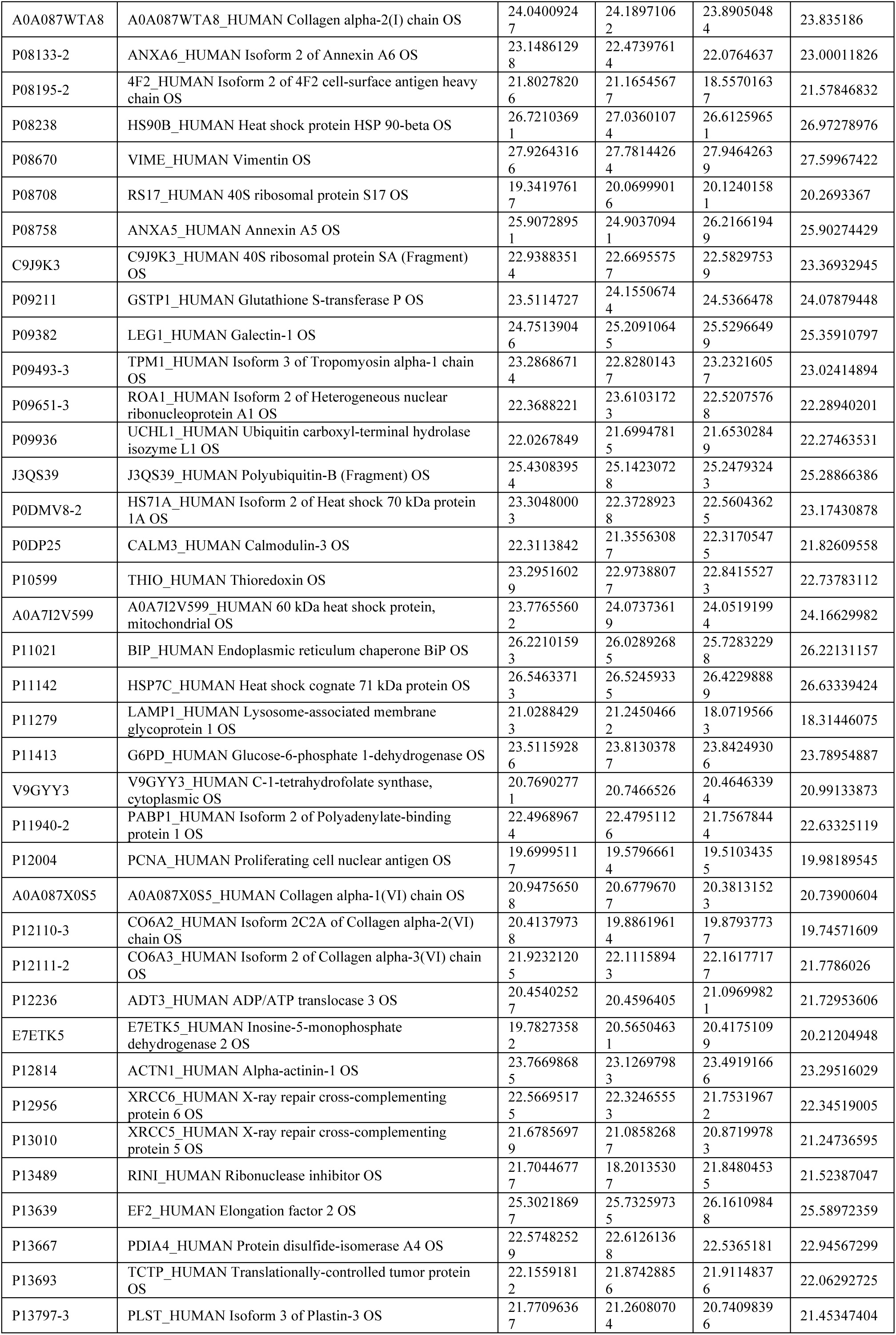
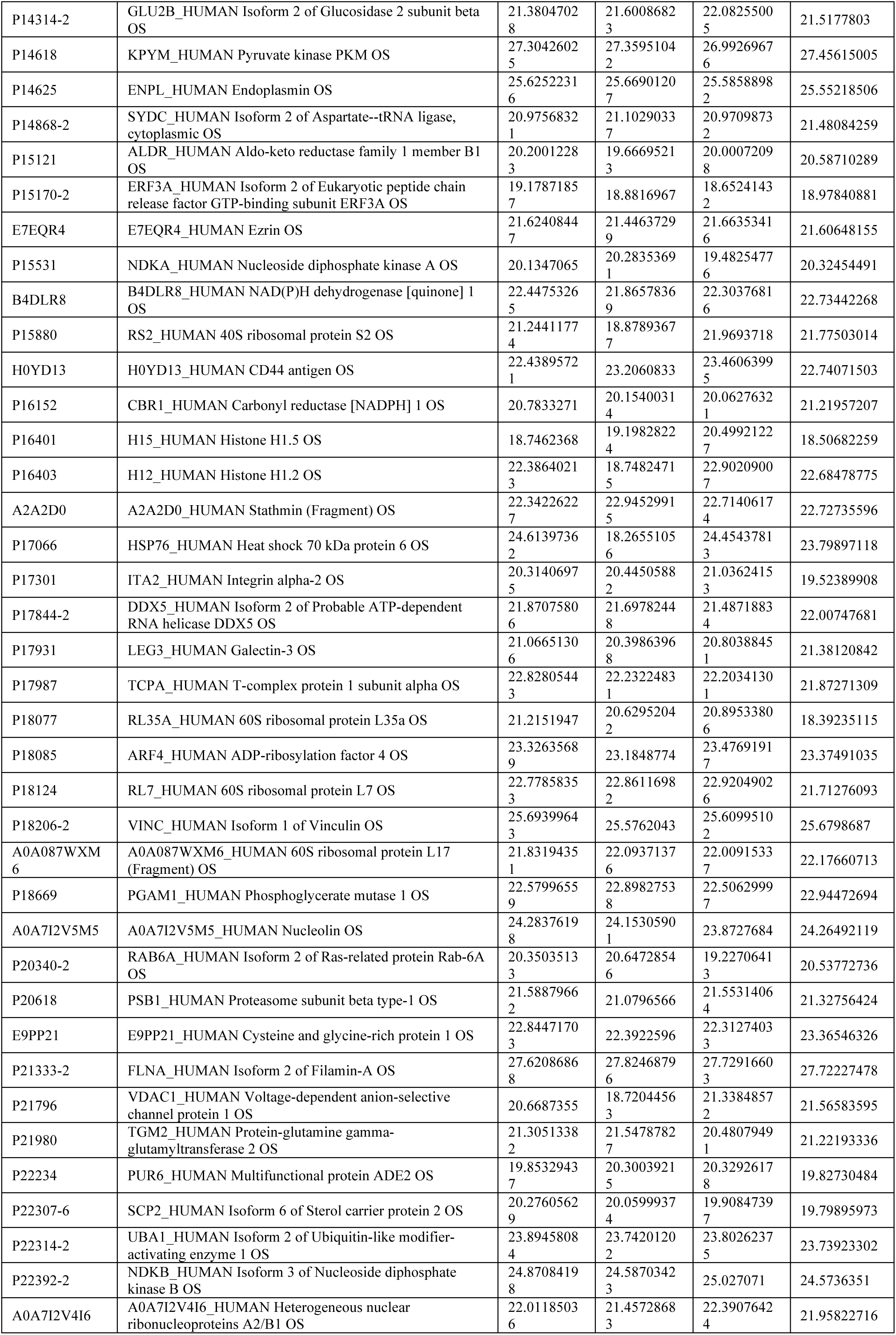
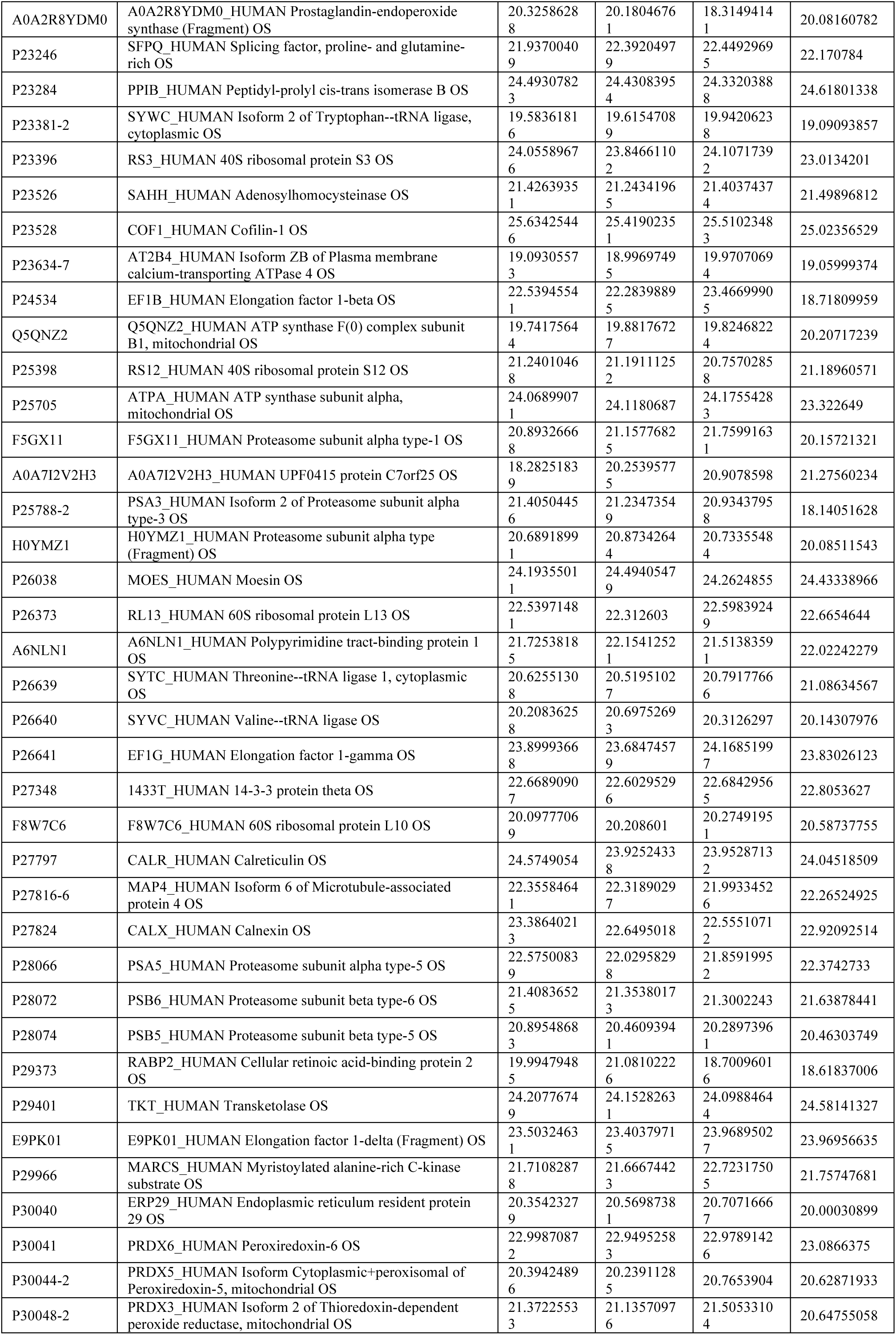
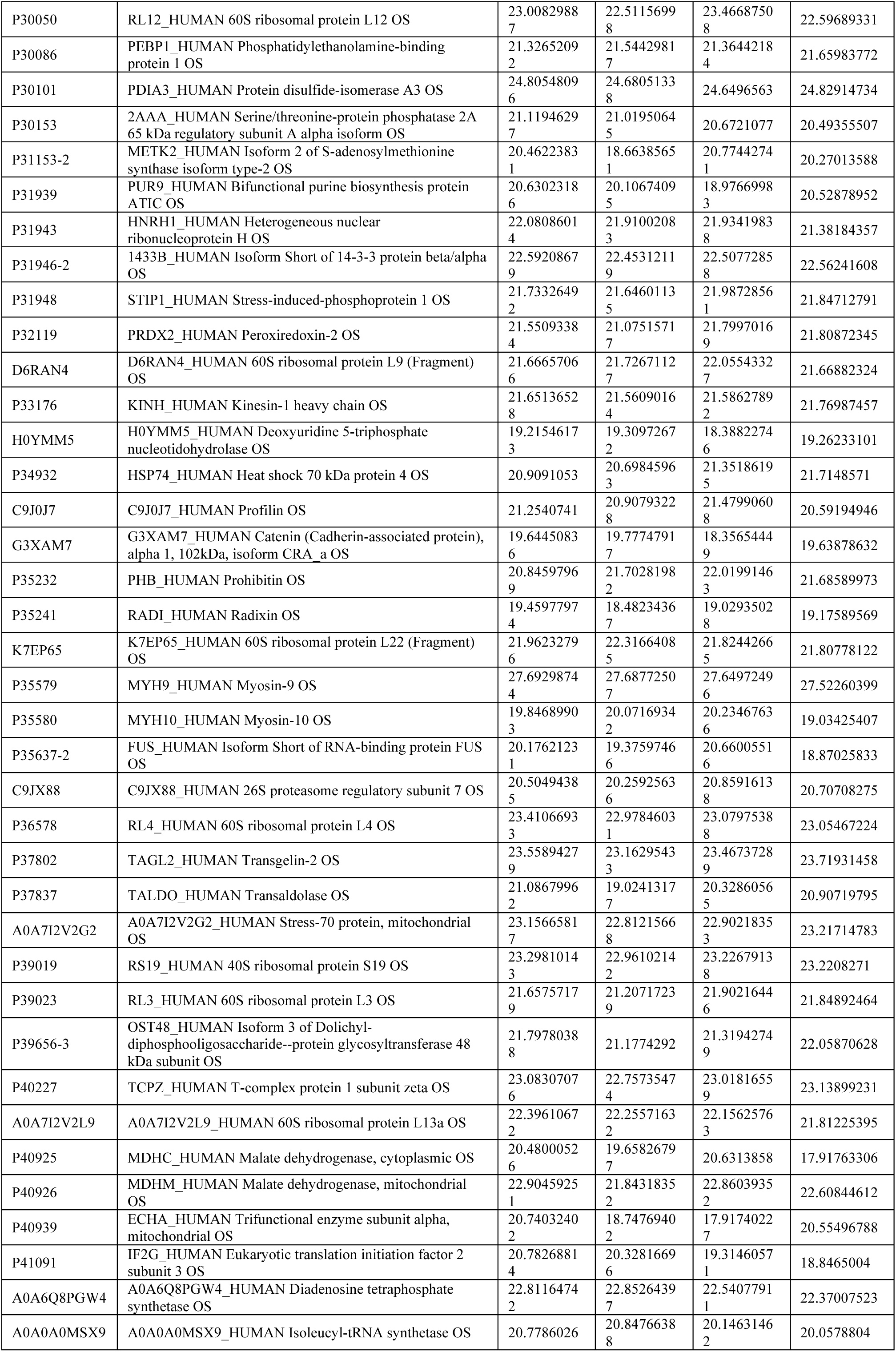
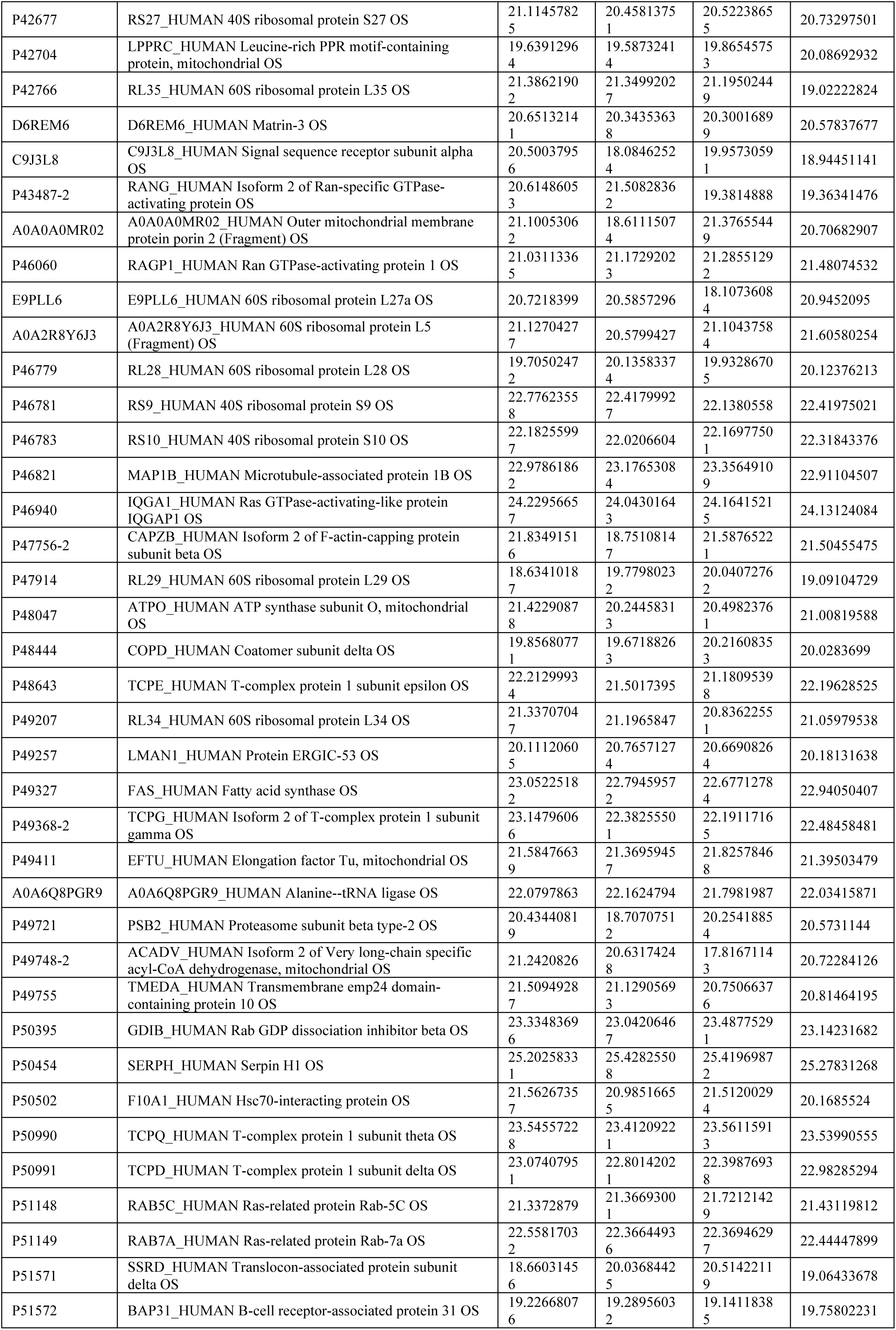
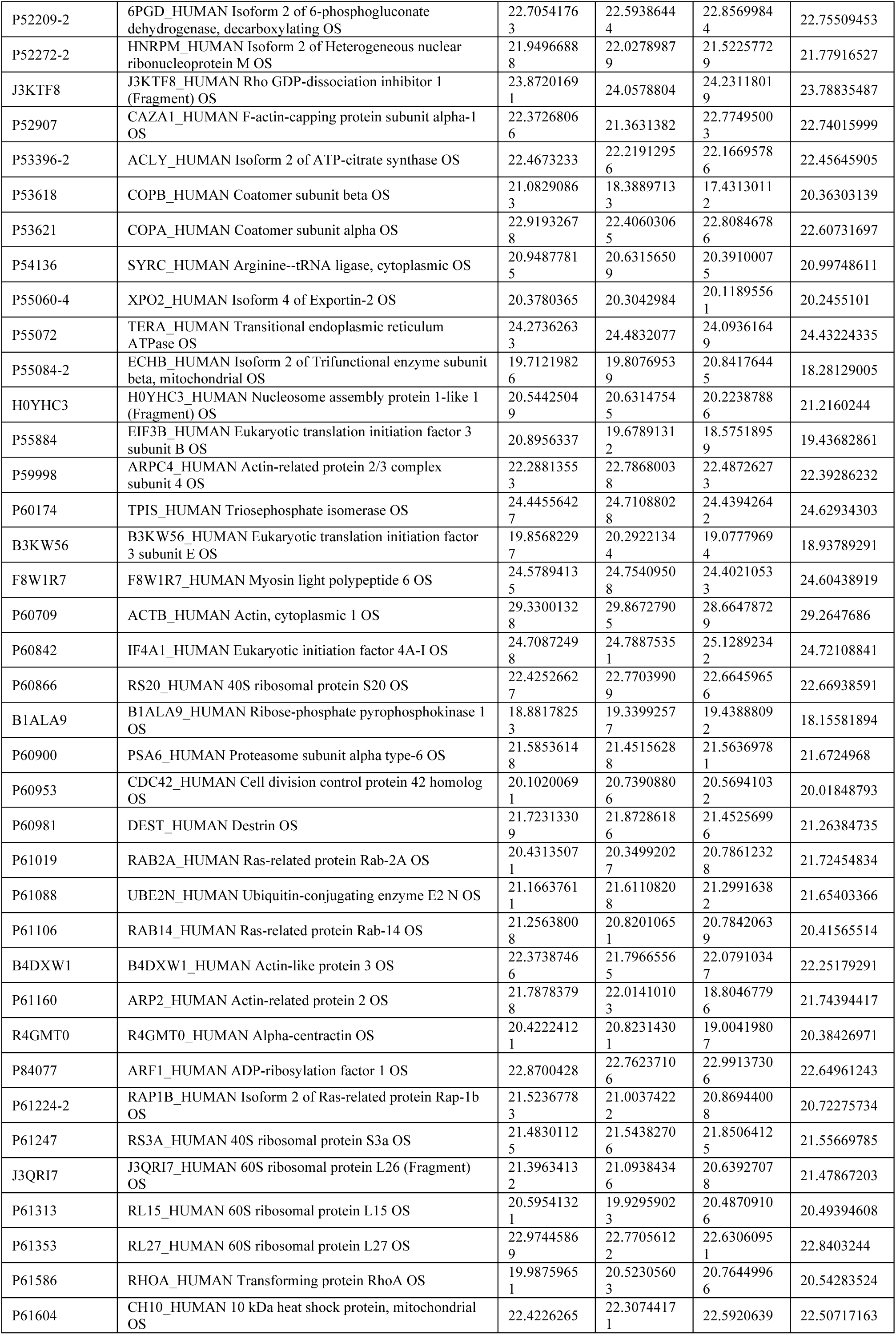
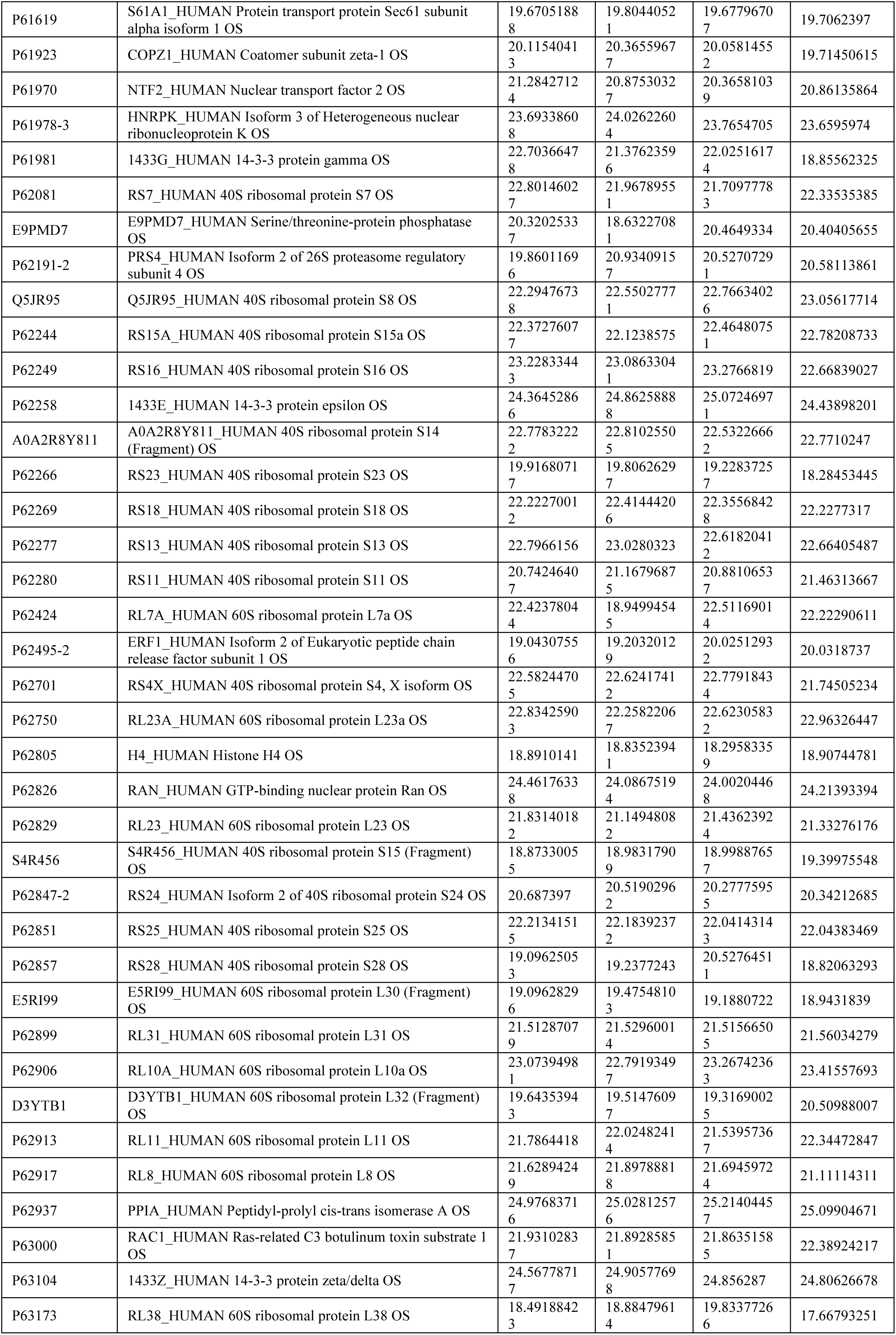
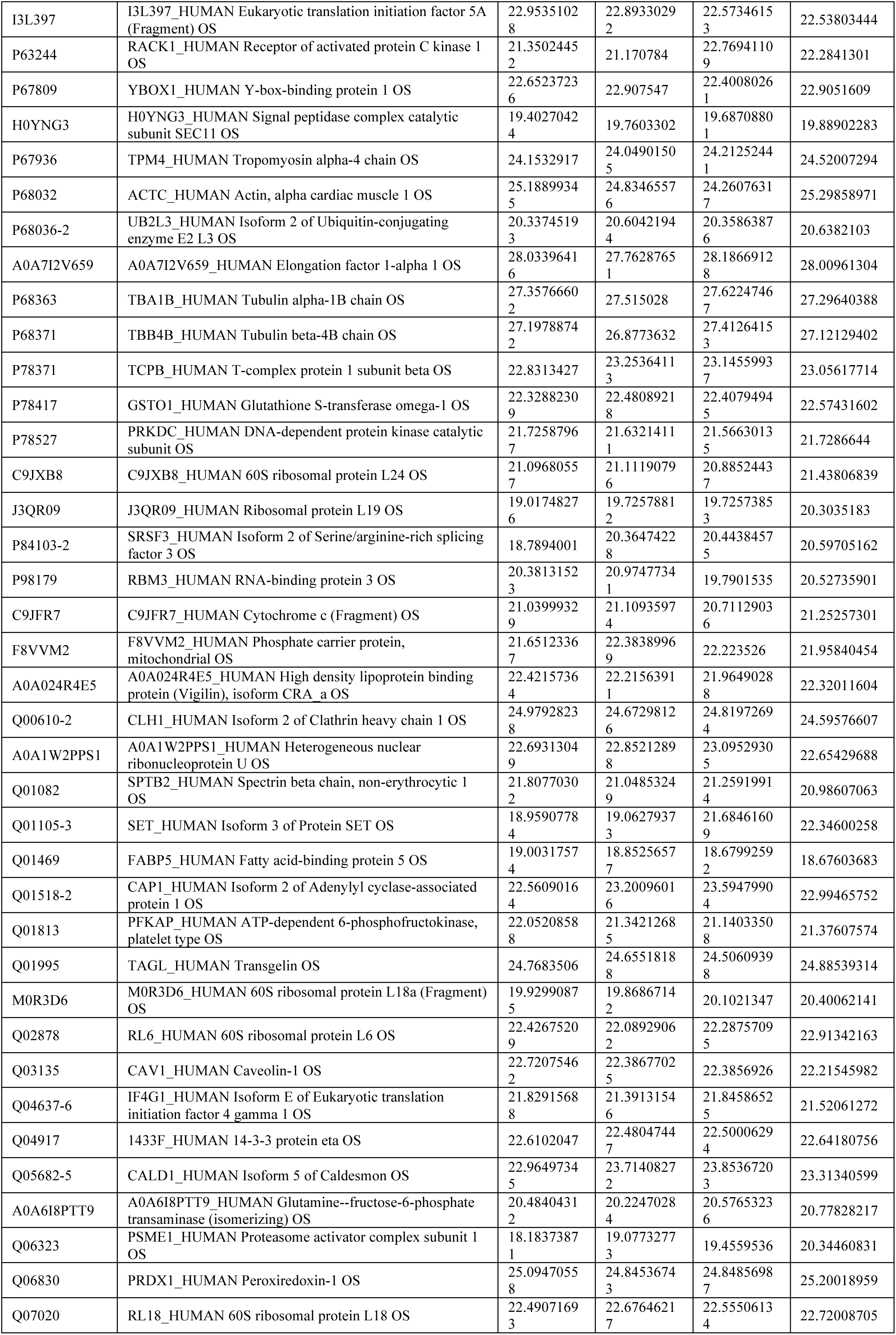
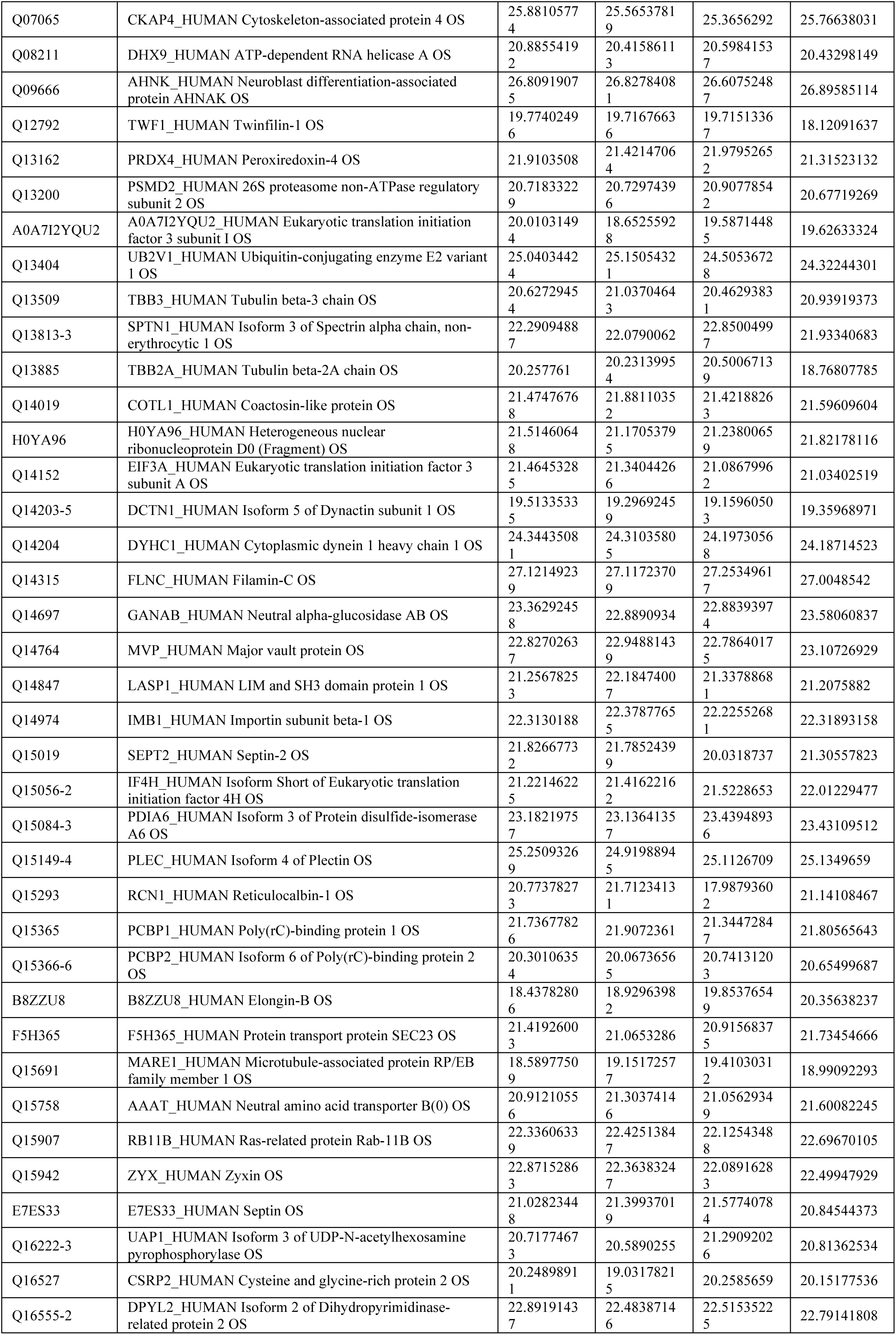
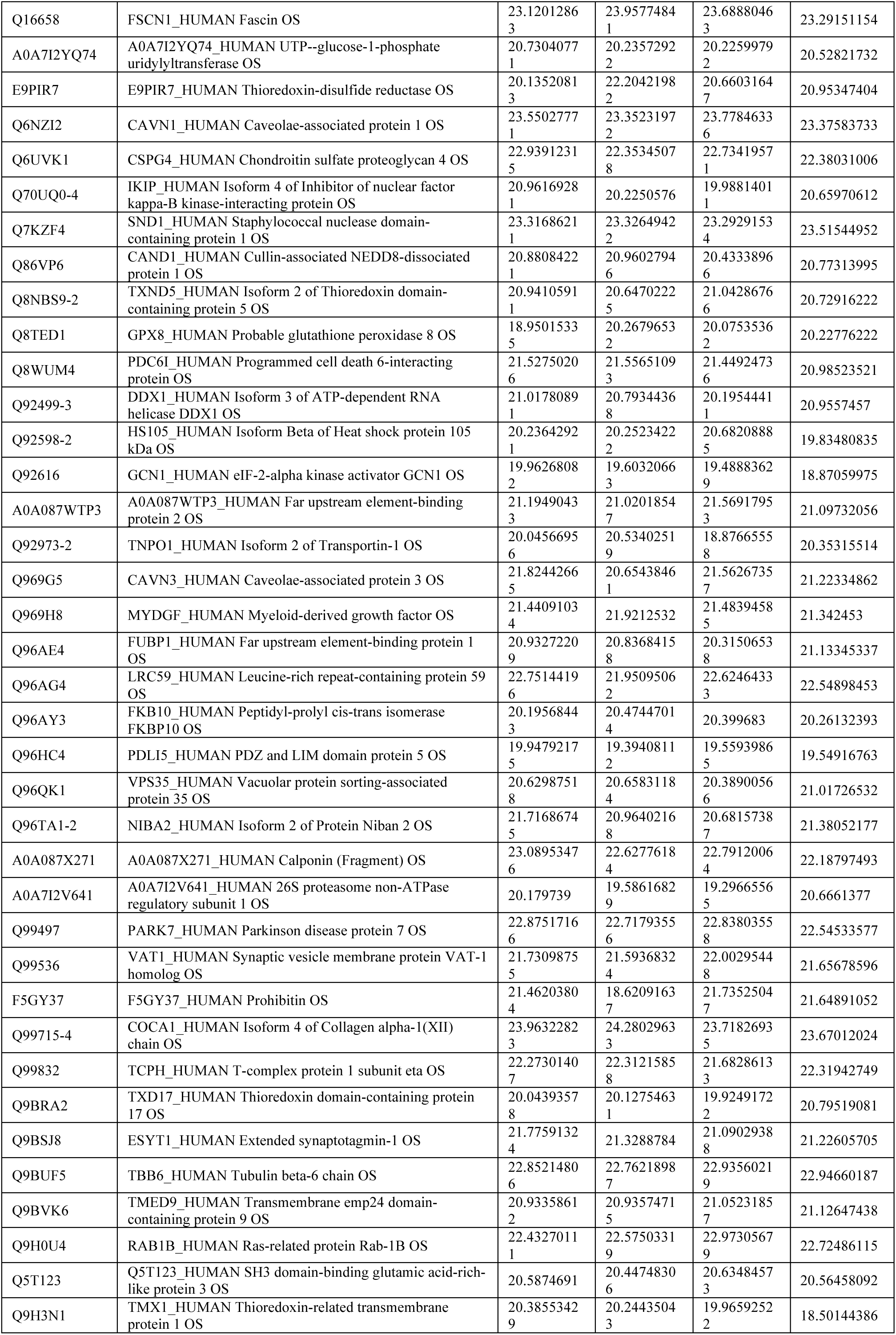
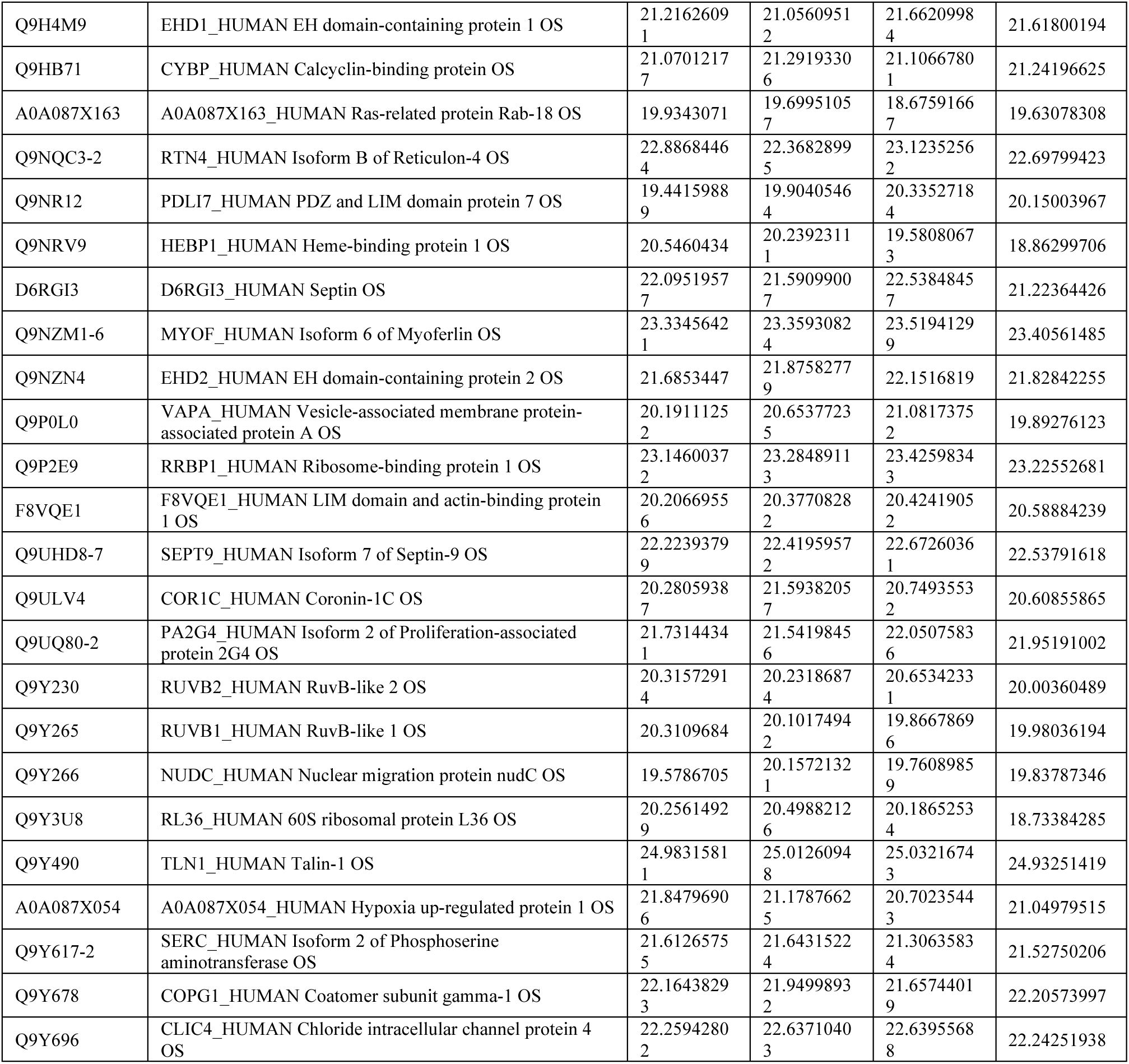
The original proteomic analysis data of the effects of SARS-CoV-2 structural proteins, as showed in Figure 4A.

**Appendix Table 2.**
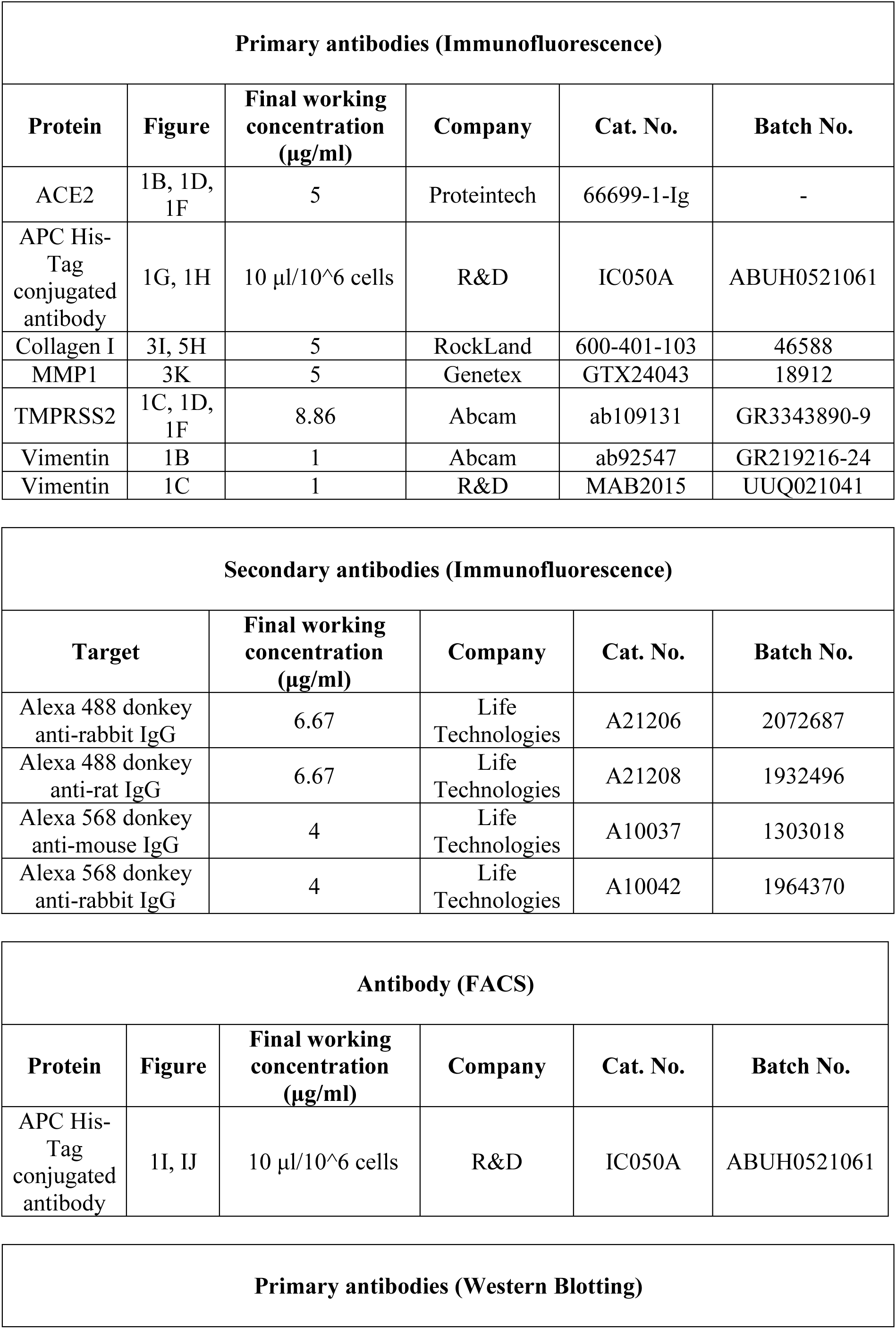
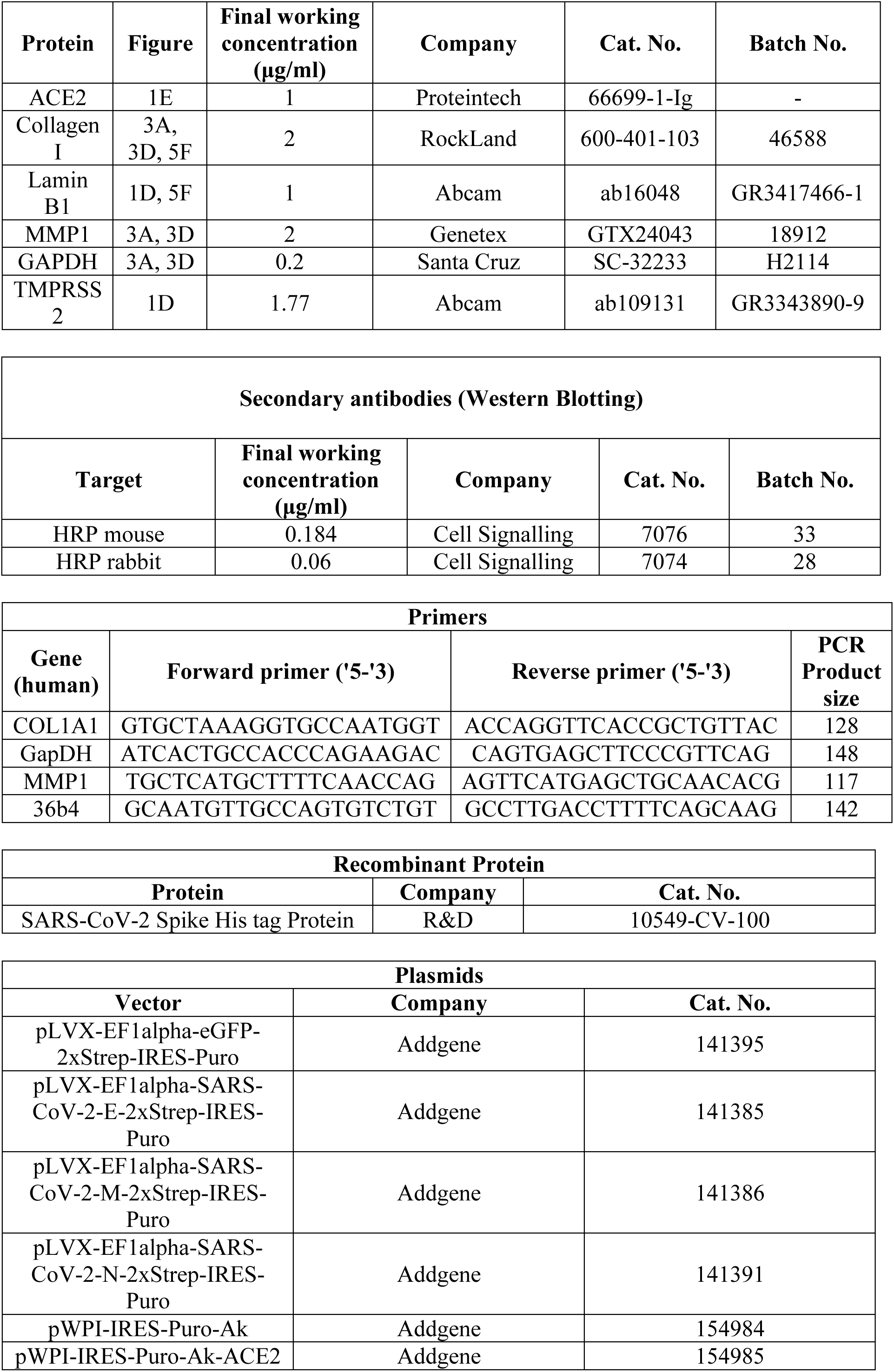
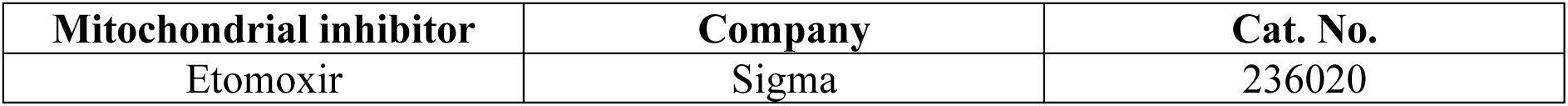
The key materials and reagents used in the current study.

